# Single cell transcriptomics reveals lineage trajectory of the retinal ganglion cells in wild-type and *Atoh7*-null retinas

**DOI:** 10.1101/2020.02.26.966093

**Authors:** Fuguo Wu, Jonathan E. Bard, Julien Kann, Donald Yergeau, Darshan Sapkota, Yichen Ge, Zihua Hu, Jie Wang, Tao Liu, Xiuqian Mu

## Abstract

Past studies concluded that Atoh7 is critical for the emergence of the retinal ganglion cell (RGC) lineage in the developing retina, whereas Pou4f2 and Isl1 function in RGC differentiation. Atoh7 is expressed in a subset of retinal progenitor cells (RPCs) and is considered a competence factor for the RGC fate, but the molecular properties of these RPCs have not been well characterized. In this study, we first used conventional RNA-seq to investigate transcriptomic changes in *Atoh7*-, *Pou4f2*-, and *Isl1*-null retinas at embryonic (E) day 14.5 and identified the differentially expressed genes (DEGs), which expanded our understanding of the scope of downstream events. We then performed single cell RNA-seq (scRNA-seq) on E13.5 and E17.5 wild-type and *Atoh7*-null retinal cells. Clustering analysis not only correctly identified known cell types at these developmental stages but also revealed a transitional cell state which was marked by *Atoh7* and genes for other lineages in a highly overlapping fashion and shared by all early developmental trajectories. Further, analysis of the *Atoh7*-null retina revealed that, unlike previously believed, the RGC lineage still progressed considerably and a substantial amount of RGC-specific gene expression still occurred. Thus, Atoh7 likely collaborates with other factors to shepherd the transitional RPCs to the RGC lineage by competing with other lineage factors and activating RGC-specific genes. This study thus provides significant insights into the nature of RPC competence for different retinal cell fates and revises our current view on the emergence of the RGC lineage.

## Introduction

The central nervous system has the most diverse cellular composition in the animal body. How this complexity is achieved during development has been one of the central questions of neuroscience. In the central nervous system, all neural cell types originate from a common pool of neural progenitor cells; neural progenitor cells take on different developmental trajectories to eventually assume distinct cell fates. The neural retina is an ideal model for studying neural development. All retinal cell types arise from a single population of retinal progenitor cells (RPCs) through a conserved temporal sequence, but with significant overlaps ^1–4^. The competence of RPCs for different cell fates change over the course of development so that different cell types are produced at different time windows ^4–12^. Several factors influencing the temporal change of RPC competence have been identified ^13–16^. However, multiple retinal cell types are often born in overlapping time windows, but the nature of RPC competence for individual cell types remains unknown. Based on gene expression, considerable heterogeneity has been observed within the general RPC population, which may be related to their competence for the various retinal cell fates ^17–24^. In agreement with this idea, many key regulators, mostly transcription factors, expressed in subsets of RPCs have been shown to regulate different retinal cell fates ^21,21,23,25–30^. Several RPC subpopulations including those expressing *Atoh7, Olig2, Neurog2*, and *Ascl1* are essential or biased for certain fates ^17,19,31,32^. However, the cell states in which these key transcription factors operate in, the actual complexity of RPC heterogeneity, the relationships between the different RPC sub-populations, and their relevance to RPC competence for individual retinal fates, have only begun to be addressed. Conventional experimental approaches have provided much insight into the genetic pathways and mechanisms underlying the formation of various retinal cell types ^1,33–35^. However, the traditional approach of investigating individual genes and cell types has been painstakingly slow-paced and inefficient in providing a comprehensive picture, since the genesis of many cell types often occurs in overlapping time and space. Understanding the relationships among different progenitor subtypes and the progression of individual cell lineages is often limited by the knowledge of marker genes, the availability of proper reagents, and the low throughput and resolution and low qualitative nature of conventional multiplexing assays. Recent development in transcriptomics analysis using next-generation sequencing, particularly at single cell levels, affords powerful means to survey the complexity of cell composition and progression of cell states for individual cell lineages.

Single cell expression profiling (single cell RNA-seq, scRNA-seq) uses microfluidic devices to isolate single cells and generate barcoded cDNA libraries. The libraries are then sequenced by next-generation sequencing ^36,37^. The sequence reads can then be decoded and attributed back to specific genes in individual cells, and expression levels of individual genes within each cell can then be determined. This approach enables the expression profiles of thousands of individual cells to be analyzed, and the cells can then be grouped (clustered) based on their similarities. The groups of cells thus identified from a developing tissue can reveal the cellular complexity of the tissue and the different cell states of individual cell lineages during development. The technology has been adopted to study many developing tissues and organs including the retina ^13,36,38–45^, and can thus be used to analyze the heterogeneity of RPCs and their relationships to different retinal lineages. The current study focuses on one of the early retinal cell types, retinal ganglion cells (RGCs). Three key transcription factors, Atoh7, Pou4f2, and Isl1, function at different stages along the RGC lineage. Atoh7 functions before the RGC fate is determined and is essential, but not sufficient, for the RGC fate ^22,25,30–32,46^, whereas Pou4f2 and Isl1 function to specify the RGC fate and promote RGC differentiation ^47–51^. Atoh7 has thus been considered a competence factor. However, how the RGC lineage emerges in the global context of retinal development and what specific roles Atoh7 plays in this process is not well understood.

To further understand RGC differentiation, we first performed conventional RNA-seq on mutant E14.5 retinas of the three key transcription factors genes, *Atoh7, Pou4f2*, and *Isl1*, to characterize the global downstream events during RGC development. This allowed us to expand the scope of downstream genes from previous studies and obtain a global view on the functions of these key regulators. We then performed scRNA-seq on retinal cells from E13.5 and E17.5 wild-type and *Atoh7*-null retinas. At these stages, particularly at E13.5, four major early retinal cell types, RGCs, horizontal cells, amacrine cells, and cones, are being generated ^4^. Our analysis not only identified all these retinal cell types with unique gene signatures but also revealed their relationship to the RPC groups. Importantly, we discovered that all the early cell lineages went through a shared transitional cell state before the cell fates were committed and that this state was marked by such genes as *Atoh7, Neurog2, Neurod1*, and *Otx2* that are involved in the formation of these lineages. Analysis of the *Atoh7*-null cells revealed that the RGC trajectory was truncated as expected, with major changes in gene expression in individual cell types/states, particularly in RGCs. Unexpectedly, the RGC lineage still formed and advanced substantially, indicating that other factors are involved in establishing this lineage. These results provide novel insights into the mechanisms governing the emergence of the different retinal lineages and particularly advance our understanding of the cellular process and genetic pathways underlying the establishment of the RGC lineage.

## Material and methods

### Animals

All mice used were in the C57BL6/129 mixed genetic background. The two knockin alleles used in this study, *Atoh7^zsGreenCREERT2^* and *Pou4f2^FLAGtdTomato^*, were described in detail in a recent publication ^52^. *Atoh7^zsGreenCREERT2^* is a null allele and *Pou4f2^FLAGtdTomato^* is a wild-type allele. The other alleles including *Atoh7^lacZ^* (null), *Pou4f2^Gfp^* (null), the conditional *Isl1*-null mice (*Isl1^flox/flox^;Six3-Cre*), and the *Atoh7^HA^* allele were reported before ^18,30,49,53^. All procedures using mice conform to the U.S. Public Health Service Policy on Humane Care and Use of Laboratory Animals and were approved by the Institutional Animal Care and Use Committees of Roswell Comprehensive Cancer Center and University at Buffalo.

### Conventional RNA-seq

Conventional RNA-seq was carried out as previously described ^29^. After timed mating, E14.5 retinas were dissected and stored in RNAlater (Invitrogen) while genotyping was performed. Three individual pools of four to six retinas were collected for individual genotypes, including wild-type, *Atoh7*-null (*Atoh7^lacZ/lacZ^*), *Pou4f2*-null (*Pou4f2^Gfp/Gfp^*), and *Isl1*-null (*Isl1^flox/flox^;Six3-Cre*). Total RNA was then isolated and RNA-seq libraries were generated using TruSeq RNA Sample Prep Kit v2 kit (Illumina, RS-122-2001) following the manufacturer’s instruction and sequenced on an Illumina HiSeq2500 sequencer. Sequence reads were mapped to the mouse genome assembly (mm10) by STAR ^54^ and differentially expressed genes (DEGs) were identified by EdgeR ^55^. The FDR cutoff was set at 0.05 and the minimum fold change imposed was 1.5. To compare gene expression changes in the three mutants, we calculated the z-scores of sequence read counts per million (CPM) for each gene, then divided the genes into five groups based on the hierarchical clustering. We then generated a heatmap of differential genes by the “pheatmap” R package (https://cran.r-project.org/web/packages/pheatmap/index.html). All RNA-seq sequence reads were deposited into the NCBI Short Read Archive (accession numbers SAMN02614558-SAMN02614569).

### Retinal cell dissociation, FACS, scRNA-seq library construction, and sequencing

Dissociation of embryonic retinas into single cell suspensions was performed as previously described ^56^. E13.5 retinas with the desired genotypes were collected after timed mating. The genotypes used in this study included *Atoh7^zsGreenCreERT2/+^* (designated as wild-type) and *Atoh7^zsGreenCreERT2/lacZ^* (*Atoh7*-null). They were then washed with cold phosphate buffered solution, pH7.0 (PBS), and transferred to fresh tubes containing 200 μl 10 mg/ml trypsin in PBS. The retinas were then incubated in a 37°C water bath for 5 mins and triturated five times with a P1000 pipette tip. 20 μl of soybean trypsin inhibitor was then added to the tube. The cells were spun down at 500 g, washed twice with PBS, and resuspended in PBS. The cells were then loaded onto the 10X Genomics Chromium Controller to generate scRNA-seq libraries using the Chromium Single Cell 3’ Library & Gel Bead Kit v2, following the manufacturer’s instructions.

We also performed fluorescence assisted cell sorting (FACS) using relatively low gating thresholds as described to enrich *Atoh7*-expressing cells and *Pou4f2*-expressing cells from E17.5 retinas carrying the *Atoh7^zsGreenCreERT2^* and *Pou4f2^tdTomato^* alleles ^52^. Cells from *Atoh7^zsGreenCreERT2/+^* and *Pou4f2^tdTomato/+^* were designated as wild-type, and those from *Atoh7^zsGreenCreERT2/lacZ^* were *Atoh7*-null. These cells were also loaded onto the 10X Genomics Chromium Controller to generate scRNA-seq libraries.

The libraries were sequenced by an Illumina HiSeq2500 rapid run using 26×8×98 sequencing, and the reads were deposited into the NCBI Short Read Archive with a GEO accession number GSE149040.

### scRNA-Seq Analysis

The output from 10X Genomics Cellranger 2.1.1 pipeline was used as input into the R analysis package Seurat version 3.1.1. Cells with high unique molecular index counts (nUMI), high mitochondrial transcript load, and high transcript counts for red blood cell markers were filtered out from the analysis. The data was then r normalized, scaled, and explored using Seurat’s recommended workflow. Principal component analysis (PCA), Louvain clustering, and the UMAP (Uniform Manifold Approximation and Projection) were performed. Using the called clusters, cluster-to-cluster differential expression testing using the Wilcoxon Rank Sum identified unique gene markers for each cluster. Differential expression between shared wild-type and mutant clusters was assessed using the FindMarkers function of Seurat, which also utilized the Wilcoxon Rank Sum test. Cell cycle analysis used a protocol in Seurat with 70 cycle genes ^57^ (https://satijalab.org/seurat/v3.1/cell_cycle_vignette.html).

To further interrogate the RGC developmental trajectory, cells belonging to C3, C4, C5, and C6 were subset from the Seurat data object. The average expression for each DEG, for each cluster, was compared between wildtype and *Atoh7*-null mice using the pheatmap package, clustering rows by euclidean distance using the hclust algorithm, and introducing cuts to the hierarchy tree using cutree = 7 for visual clarity.

### Pseudotime Analysis

To infer developmental trajectories, the python package SCANPY provides pseudotemporal-ordering and the reconstruction of branching trajectories via Diffusion Pseudotime (DPT) ^58^. A root cell was selected at random within the progenitor cell population of called Cluster 1. The assigned pseudotime values were then mapped back to the Seurat UMAP embedding for visualization and further analysis.

### Immunofluorescence staining, In situ hybridization and online data mining

Immunofluorescence staining on cryopreserved retinal sections was performed as described before ^29,59^. Primary antibodies used in this study included: rabbit anti-Otx2 (1:200, Sigma, B74059), goat anti-Olig2 (1:200, R&D system, AF2418), mouse anti-Neurog2 (1:200, R&D system, MAB3314), goat anti-HA (1:100, Genscript, A00168), rabbit anti-Atoh7 (1:200, Novus, NBP1-88639), rabbit anti-Uchl1 (Pgp9.5) (1:500, Millipore, AB1761), mouse anti-Nefm (1:200, Sigma, N5264). Immunofluorescence images were captured by a Leica TCS SP2 confocal microscope. Positive cells were counted manually per arbitrary length unit as previously described ^29,56,59^.

In situ hybridization was performed using RNAscope double Z probes (Advanced Cell Diagnostics) on paraffin-embedded retinal sections. After timed mating, embryos of desired stages were collected, fixed with 4% paraformaldehyde, embedded in paraffin, sectioned at 6 μm, and de-waxed, as previously described ^29,49,59,60^. The sections were then processed, hybridization was performed, and the signals were visualized using the RNAscope^®^ 2.5 HD Detection Reagents-RED following the manufacturer’s manual. In situ images were collected using a Nikon 80i Fluorescence Microscope equipped with a digital camera and Image Pro analysis software.

## Results

### Changes in gene expression in *Atoh7*-, *Pou4f2*-, and *Isl1*-null retinas

Atoh7, Pou4f2, and Isl1 are three key regulators in the gene regulation network controlling RGC development ^49,59^. They function at two different stages; Atoh7 is believed to confer competence to RPCs for the RGC lineage, whereas Pou4f2 and Isl1 function to specify the RGC fate and promote differentiation. Previously, downstream genes of Atoh7, Pou4f2, and Isl1 have been identified by microarrays ^22,49,60,61^. However, due to limitations of the technology used, those genes likely only cover small proportions of those regulated by the three transcription factors. To gain a more global view of the function of the three transcription factors, we collected total RNA samples from wild-type, *Atoh7*-null, *Pou4f2*-null, and *Isl1*-null retinal tissues at E14.5 and performed RNA-seq. The RNA-seq data from the wild-type retina provided a comprehensive list of genes expressed in the E14.5 retina with at least 1.0 average counts per million reads (CPM, see Suppl. Table 1). We then identified differentially expressed genes (DEGs) by edgeR ^55^ in each of these mutant retinas as compared to the wild-type retina using a cutoff of at least 1.5 fold change and FDR of at least 0.05 (Suppl. Tables 2-4). In the *Atoh7*-null retina, we identified 670 downregulated genes including *Pou4f2* and *Isl1*, and 293 upregulated genes (Suppl. Table 2); in the *Pou4f2*- mull retina, we identified 258 downregulated genes and 169 upregulated genes (Suppl. Table 3); and in the *Isl1*-null retina, we identified 129 downregulated genes and 79 upregulated genes (Suppl. Table 4). Although *Atoh7* and *Pou4f2* were also identified as DEGs in their own respective mutant retinas, *Isl1* showed no change in the *Isl1*-null retina, because in the *Isl1*-null retina only the small frame-shifting exon 3 was deleted, which likely did not alter the mutant mRNA levels significantly ^49^. These DEGs not only confirmed previous findings, as essentially all previously identified DEGs were included, but also provided a more complete picture by significantly increasing the numbers of DEGs for each mutant. The different numbers of DEGs in these three mutant retinas were consistent with their severity of defects in RGC development ^25,30,49,50,53^, which was further reflected by a clustering heatmap analysis, showing that the *Atoh7*-null retina was least similar, and the Isl1-null retina was most similar, to the wild-type retina (Figure 1a).

**Figure 1.**
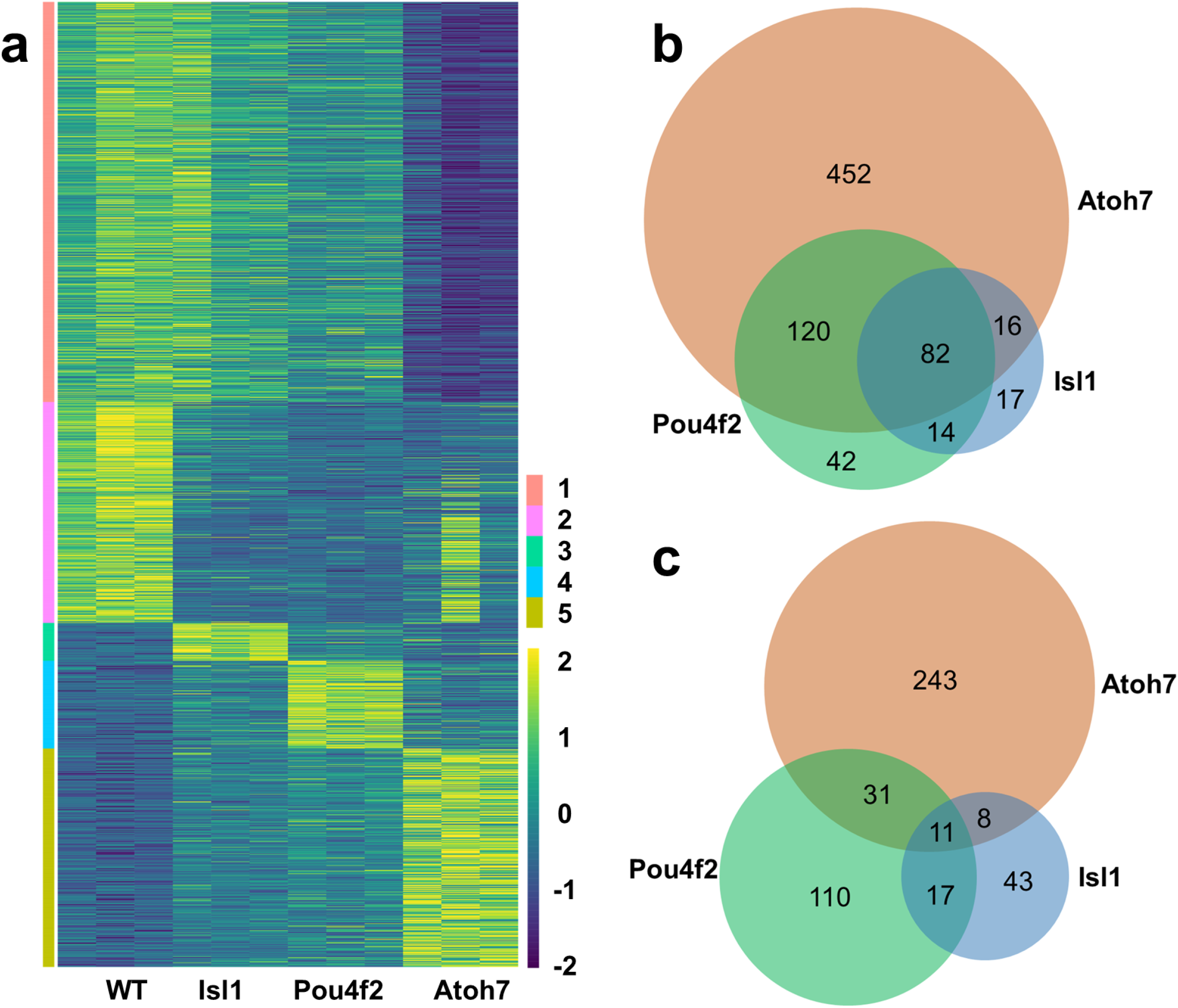
Conventional RNA-seq identifies differentially expressed genes (DEGs) in E14.5 *Atoh7*-null, *Pou4f2*-null, and *Isl1*-null retinas. **a**. Clustering of all DEGs across all four genotypes based on Z scores indicates the *Atoh7*-null retina is least similar, whereas the *Isl1*-null retina is most similar to the wild-type retina. The genes were divided into five groups (1-5) based on how they are affected in the three mutants. **b**. A Venn-diagram showing the overlaps of downregulated genes in all three mutant retinas. **c**. A Venn-diagram showing the overlaps of upregulated genes in the three mutant retinas.

Consistent with RGCs being largely missing in the *Atoh7*-null retina, RGC-specific genes were mostly found in the downregulated DEG list, whereas RPC-expressed DEGs included both down- and upregulated genes (Suppl. Table 2). The downregulated Atoh7 gene list also encompassed the majority of genes in downregulated Pou4f2 and Isl1 lists, as would be expected considering that Pou4f2 and Isl1 function downstream of Atoh7 (Figure 1b). Gene ontology (GO) analysis by DAVID ^62^ found the downregulated genes in the three mutants were highly associated with different aspects of neural differentiation, as demonstrated by enriched biological processes and GO terms, including nervous system development, axonogenesis, cell adhesion, and neurotransmitter secretion (Suppl. Table 5). On the other hand, the three upregulated gene lists were markedly different and much less overlapped (Figure 1c, Suppl. Tables 2-4). GO analysis revealed the upregulated DEGs in the three mutants were broadly involved in neural development, both negatively and positively (Suppl. Table 5). Genes negatively regulating proliferation were enriched in the Atoh7 upregulated list, but not the Pou4f2 or Isl1 upregulated list (Suppl. Table 5). These results reflected that these three factors repress gene expression largely independently at two different levels of the gene regulatory hierarchy either directly or indirectly; Atoh7 represses gene expression in proliferating RPCs, confirming our previous analysis ^22^, whereas *Pou4f2* and *Isl1* repress gene expression in RGCs. Independent gene repression by these factors was also demonstrated by genes with changes in different directions in these three mutant retinas (Suppl. Tables 2-4). For example, *Nhlh1* did not change in *Atoh7*-null, was down-regulated in *Pou4f2*-null (fold change −2.2), but upregulated in *Isl1*-null (fold change 1.7), whereas its related gene *Nhlh2* was down-regulated (fold change −2.0) in *Atoh7*-null, did not change in *Pou4f2*-null, but was significantly up-regulated in *Isl1*-null (fold change 1.7). We also confirmed that some marker genes for amacrine cells (e.g. *Chat, Th*, fold change 383.0 and 35.1 respectively) were markedly up-regulated in *Pou4f2*-null as previously reported ^61^, but they did not change in either *Atoh7*-null or *Isl1*-null retinas. *Dlx1* and *Dlx2*, two genes involved in RGC development ^63,64^, were down-regulated in the *Atoh7*-null retina, but up-regulated in the *Pou4f2*- and *Isl1*-null retinas, indicating these genes were activated by Atoh7 in RPCs, but repressed by Pou4f2 and Isl1 in RGCs (Suppl. Tables 2-4).

The DEG lists also revealed/confirmed that key pathways were affected in the three mutant retinas and additional components were found to be affected. For example, the Shh pathway, which is under the control of the gene regulatory network for RGC genesis, plays a key role in balancing proliferation and differentiation through a feedback mechanism ^49,50,60,65–68^. Expanding previous findings, we found more components in the Shh pathway were affected in all three mutant retinas (Figure 2a, Suppl. Tables 2-4). These component genes, including *Shh, Gli1, Ptch1, Ptch2*, and *Hhip*, revealed a complex feedback loop of the pathway in balancing proliferation and differentiation via downstream genes such as *Gli1* and *CcnD1* (Cyclin D1) (Suppl. Tables 2-4, Figure 2b) ^49,50,60,65–71^. Consistent with this model, expression of *Gli1* and *CcnD1* was reduced in all three mutants (Suppl. Tables 2-4). Two additional signaling pathways from RGCs to RPCs exist in the developing retina. Two related BMP molecules, Gdf11 and Myostatin/Gdf8 (Mstn), are also secreted from RGCs to balance RGC production and RPC proliferation ^60,72^. Vegf is also involved in the feedback from RGCs to RPCs ^73^. *Mstn* was downregulated in all three mutant retinas, but *Gdf11* exhibited no change, which may due to *Gdf11* expression not confined to just RGCs (data not shown). Unexpectedly, *Vegfa* expression increased in the Atoh7-null retina, although the underlying mechanism is not clear, but did not change in the other two mutants. Multiple component genes of the Notch pathway were upregulated in the *Atoh7*-null, but not the other two mutant retinas (Suppl. Tables 2-4). Some genes in the Notch pathway such as *Hes5* were also affected in the Atoh7-null retina (Suppl. Tables 2). However, as discussed later with our single cell analysis, the effects of *Atoh7* deletion on the Notch pathway is complex and likely involved the Shh and Vegf pathways, as crosstalk exists between them via *Hes1* ^66,68,73,74^. These results further demonstrated that deletion of *Atoh7* not only compromised RGC formation but also altered the properties of RPCs through multiple interacting pathways.

**Figure 2.**
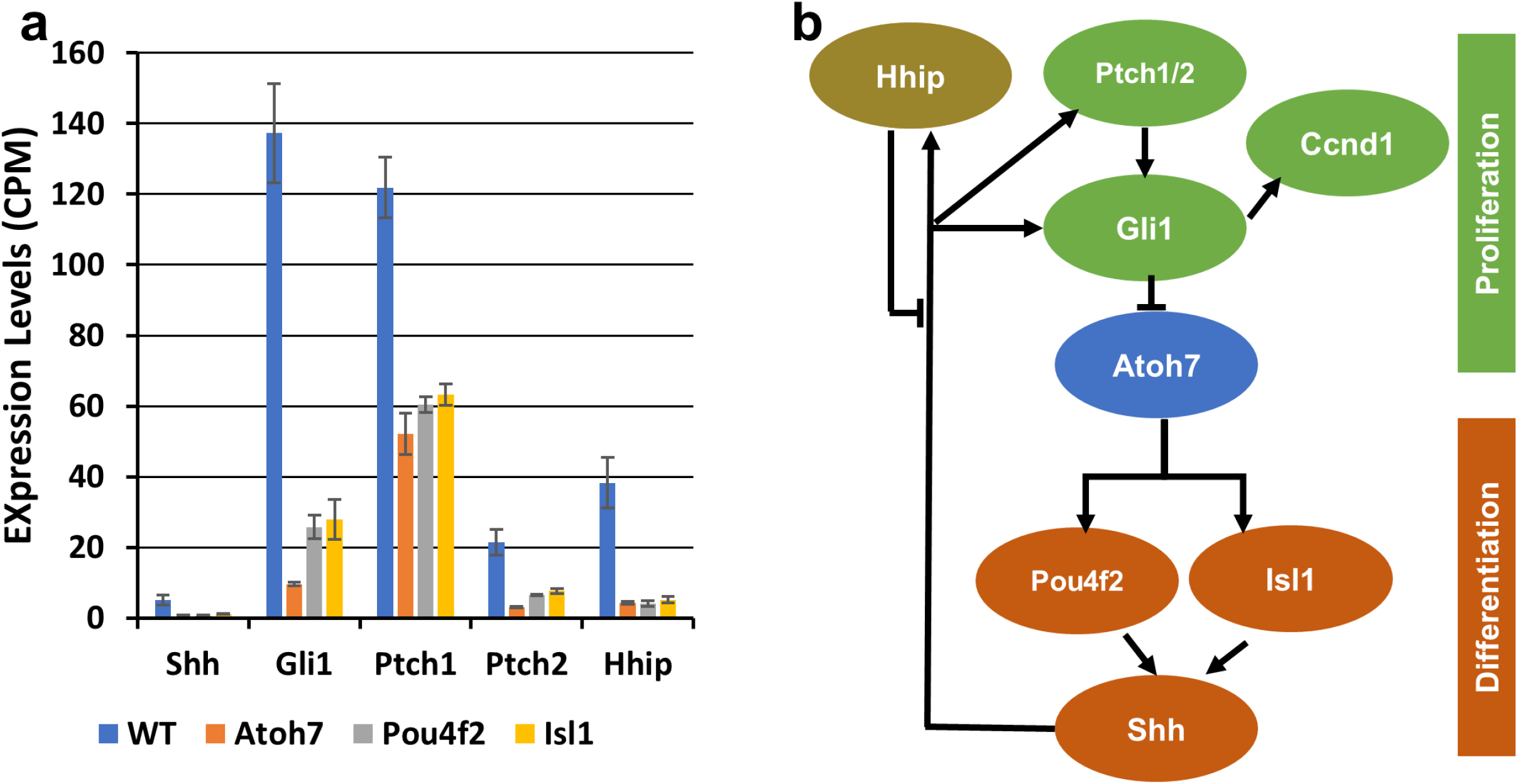
The Shh pathway is regulated by the RGC gene regulatory cascade at multiple levels. **a**. Multiple component genes of the Shh pathway are downregulated in the three mutant retinas. Error bars are standard deviation. P values between all mutants and wild-type were all smaller than 0.001. **b**. A diagram showing how the *Atoh7*-*Pou4f2/Isl1* gene regulatory cascade regulates the Shh pathway in the developing retina in a complex manner, with multiple feedback loops.

### Single cell RNA-seq of wild-type and *Atoh7*-null retinas at E13.5

Whereas the RNA-seq data provided much insight into gene regulation by Atoh7, Pou4f2, and Isl1 in the developing retina, how their absence affected different cell states could not be attained. Particularly for Atoh7, it functions in a subset of RPCs that gives rise to RGCs, but the properties of these RPCs and their relationships to other cell populations have not been well characterized. To that end, we first performed single cell RNA-seq with E13.5 wild-type and *Atoh7*-null retinal cells. The choice of E13.5, instead of E14.5, was fortuitous but did not affect our overall analysis since largely the same cell types are being generated in these two time points ^2,4^. After filtering out blood cells, doublet cells, and stressed cells, we were able to obtain expression data of 3521 wild-type cells and 6534 *Atoh7*-null cells. The median sequence reads were 68,491 and 54,765 for wild-type and mutant cells respectively. The median numbers of genes captured were 1,975 and 2,375 for wild-type and mutant cells respectively. UMAP clustering was then performed on these cells using Seurat 3.1.1 ^75^, which resulted in a total of 11 clusters (C0-C10) for both wild-type and *Atoh7*-null cells, and the corresponding clusters highly overlapped (Figure 3a, b). We first used known marker genes to assign identities to these clusters. These markers included *Ccnd1, Fgf15*, and *Sox2* for naïve RPCs ^24,65,76^, *Sox2, Atoh7* and *Otx2* for subpopulations of RPCs ^18,23,77^, *Pou4f2* and *Pou6f2* for RGCs ^78,79^, *Ptf1a* and *Tfap2b* for amacrine and horizontal precursor cells ^27,28^, *NeuroD4* and *Crx* for photoreceptors ^22,80,81^, and *Otx1* and *Gja1* for ciliary margin cells ^82,83^. At this stage, horizontal cells and amacrine cells seemed not to have fully diverged yet and thus were grouped together (Figure 3a, b). These marker genes were specifically expressed in distinct clusters as demonstrated by dot plots (Figure 3c) and feature plot heatmaps (Suppl. Figure 1). This allowed us to definitively designate their identities, including three clusters as naïve RPCs (C0-C2), two as transitional RPCs (C3 and C4) for reasons further discussed below, two as RGCs (C5 and C6), one as horizontal and amacrine precursors (C7), two as photoreceptors (cones) (C8 and C9), and one as ciliary margin cells (C10). Notably and as expected, *Atoh7* was absent and the two RGC markers *Pou4f2* and *Pou6f2* were markedly diminished in the *Atoh7*-null cells, but the corresponding clusters in which they were expressed in the wild-type, including the two transitional RPC clusters (C3, C4) and two RGC clusters (C5, C6), still existed (Figure 3b, c, Suppl. Figure 1). Marker genes for the other mutant clusters did not show overt changes in their expression (Figure 3c, Suppl. Figure 1). These results were consistent with previous knowledge that RGCs, horizontal cells, amacrine cells, and cones are the major cell types being generated at this developmental stage ^2,4^, and that deletion of *Atoh7* specifically affects RGCs ^25,30^.

**Figure 3.**
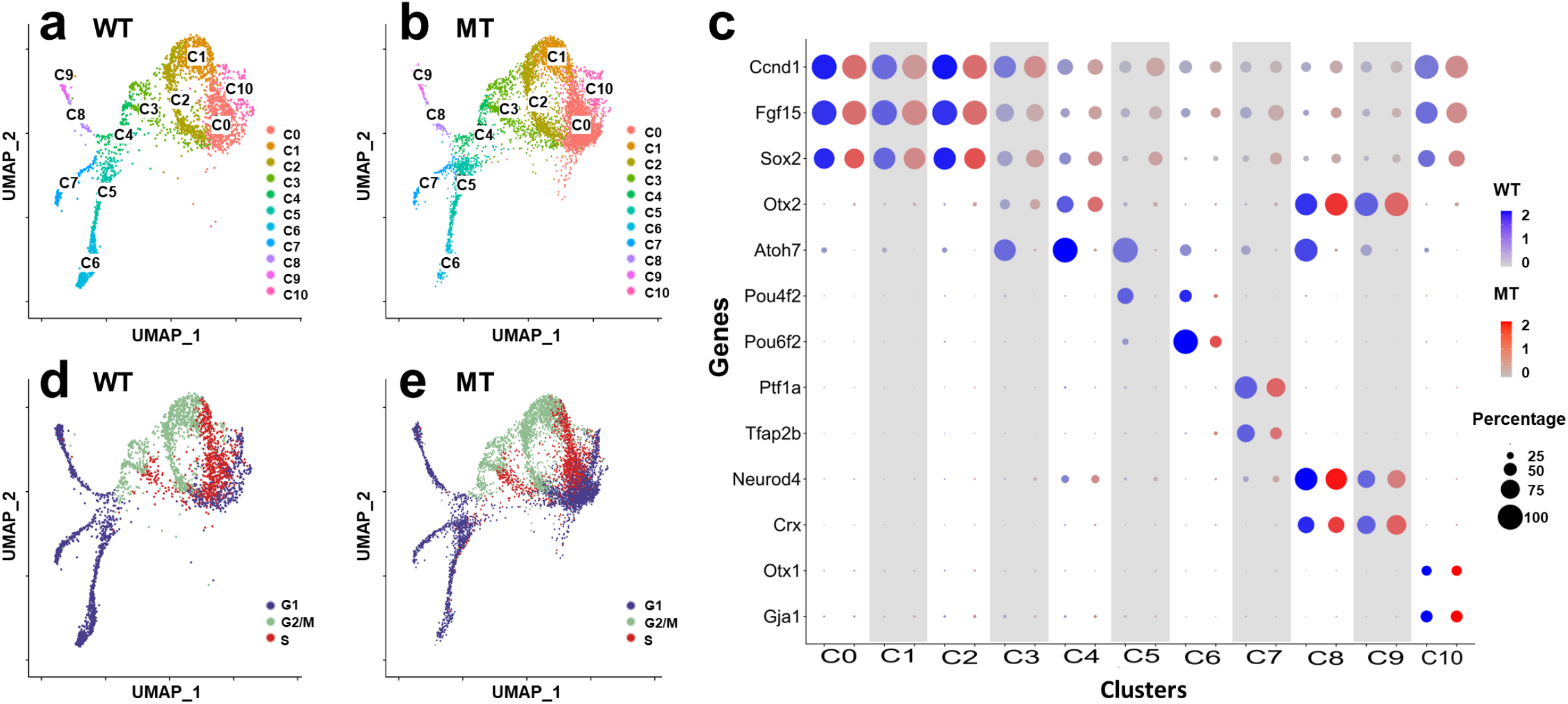
scRNA-seq analysis of E13.5 wild-type (WT) and *Atoh7*-null (MT) retinas. **a**. **b**. As indicated, UMAP clustering leads to the same 11 overlapping clusters (C0-C10) with both WT (**a**) and MT (**b**) retinal cells. **c**. Expression of known markers genes as represented by a dot plot enables identity assignment of individual WT and MT clusters (see text for details). As indicated, the sizes of the dots indicate the percentage of cells expressing the gene in individual clusters and the color intensities denote average expression levels. **d**. **e**. Cell cycle analysis determines the cell cycle status (G1, S, and G2/M) of individual cells in the clusters. Note that largely the same cell cycle distribution is observed in the WT (**d**) and MT (**e**) retinas, and that the cell cycle statuses correlate with UMAP clustering.

We also performed cell cycle analysis following a protocol in Seurat 3 using 70 cell cycle markers ^57^ and found that, for both wild-type and *Atoh7*-null cells, the three naïve progenitor cell clusters C0-C2 roughly co-segregated with their positions in the cell cycle, C0 in G1 and early S, C1 in S and G2/M, and C2 in G2/M (Figure 3d, e). The transitional RPCs (C3, C4) were also actively proliferating as they were found in different phases of the cell cycle, C3 in S and G2/M and C4 in G1. On the other hand, clusters composed of differentiating cells (C5, C6, C7, C8, C9) were all in G1 (G0) phase, confirming that they were indeed postmitotic neurons. These results demonstrated that our clustering analysis accurately grouped the cells into different stages of differentiation and our identity assignments were accurate.

### Relationships between the clusters

To further examine the characteristics of the individual clusters, we performed gene enrichment analysis of the wild-type cells by comparing the expression profile of each cluster with those of all the other clusters and identified genes that were specifically enriched in individual clusters (Suppl. Table 6). The numbers of enriched genes in these clusters ranged from 126 to 686 with Cluster 6 having the most enriched genes (Suppl. Tables 6 and 7). The enriched genes further confirmed our initial cluster identity assignment, as many additional known marker genes specific for the cell states/types were enriched in the corresponding clusters (Suppl. Table 6). Examples of such genes included *Sfrp2, Lhx2, Zfp36l1*, and *Vsx2* for RPCs (C0-C2) ^14,84–86^, *Isl1, Nefl, Sncg, Gap43*, and *Ina* for RGCs (C5 and C6) ^22,49,60^, *Thrb, Meis2, Prdm1*, and *Gngt2* for photoreceptors (C8 and C9) ^87–90^, *Tfap2a, Prdm13*, and *Onecut2* for amacrine and horizontal cell precursors (C7) ^29,91,92^, and *Ccnd2* and *Msx1* for ciliary margin cells (C10) ^93,93,94^.

Next, we examined the expression of the top ten enriched genes as ranked by p values from each wild-type cluster across all the clusters and represented the data by a heatmap (Figure 4a). This analysis did not only confirm their enrichment in the corresponding clusters but also revealed that many of these genes were expressed across several neighboring clusters, suggesting the relationships and continuity among these clusters along different developmental lineages. For example, the top ten enriched genes in C0 were also highly expressed in C1 and C2, indicating they indeed were all RPC clusters. The differences among these three clusters were likely due to their cell cycle status (Figure 3d), as many of the cluster-specific genes are directly involved in cell cycle regulation (Suppl. Table 6). C3 and C4 were two other examples of this continuity. They continued to express many of RPC genes enriched in C0-C2, albeit at lower levels, but began to express such genes as *Atoh7, Dlx1, Dlx2, Neurod1*, and *Otx2* which regulate retinal cell differentiation ^23,25,30,64,95^. On the other hand, many of the genes in C3 and C4 trailed into the further differentiated clusters including C5 and C6 (RGCs), C7 (horizontal and amacrine cells), and C8 and C9 (photoreceptors), suggesting that C3 and C4 cells were intermediate transitional RPCs poised to differentiate (Figure 4a). Although C5 and C6 were both assigned as RGC clusters, C5 continued to express many genes enriched in C3 and C4, but C6 essentially stopped expressing them (Figure 4a). On the other hand, although C5 cells expressed the early RGC marker genes such as *Isl1* and *Pou4f2* at high levels, they had not or had just begun to express many of the RGC-specific genes encoding RGC structure and function proteins such as *Nefl, Sncg, Gap43, Nefm*, and *Ina*, but these genes were highly expressed C6 cells (Figure 4a). Thus, C5 cells were nascent RGCs and C6 were further differentiated RGCs. Similarly, C8 were nascent photoreceptors and C9 were more differentiated photoreceptors based on the expression of early and later photoreceptor marker genes (Figure 4a). As mentioned above, C7 cells were considered precursors for horizontal and amacrine cells as they expressed genes required for both lineages such as *Ptf1a* but more specific marker genes for the two cell types were not robustly expressed yet. From these overlaps in expression the trajectories of the different cell lineages could be postulated, which all started from the naïve RPCs (C0-C2), underwent the transitional RPC stage (C3, C4), and finally reached the different terminal cell fates (C5-C9).

**Figure 4.**
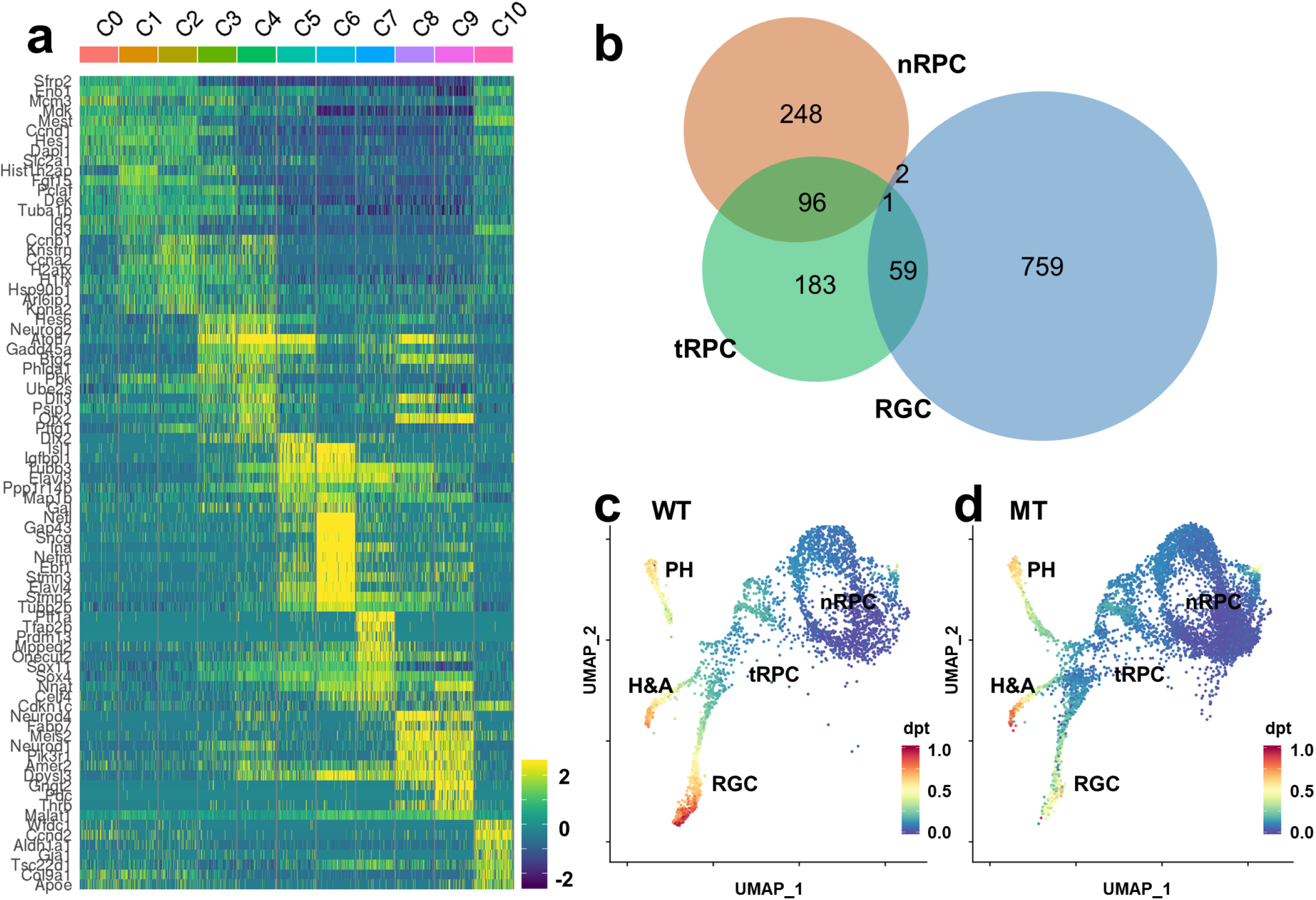
Cluster-specific gene expression reveals the relationships among the wild-type clusters. **a**. A heatmap showing the top ten enriched genes in each cluster. Each horizontal line represents one gene and each vertical line represents one cell. Data from equal numbers (100) of cells from each cluster are shown. The heatmap demonstrates the continuity and directionality between these clusters. **b**. A Venn diagram showing the overlaps of enriched genes between naïve RPCs (nRPC, C0-C2), transitional RPCs (tRPC, C3 and C4), and RGCs (C5 and C6). tRPCs have substantial overlaps with both nRPCs and RGCs, but nRPCs and RGCs have very few overlapped genes, indicating the unidirectional relationship of these clusters. **c**. **d**. Developmental trajectories predicted by the SCANPY tool based on diffusion pseudotime (DPT). Three trajectories representing the emergence of photoreceptors (PH), horizontal and amacrine cells (H&A), and RGCs from RPCs are identified for both the wild-type (WT) (**c**) and *Atoh7*-null (MT) (**d**) cells. Progression is color-coded and the direction of each lineage is clearly discernible, although the RGC lineage of the MT cells does not advance as far as the WT cells.

The unidirectional trajectories were further validated by examining the overlaps of all enriched gene lists in different clusters; there were significant overlaps between the naïve RPCs (C0-C2) and transitional RPCs (C3, C4), between the transitional RPCs (C3, C4) and the three terminal lineages including the RGC clusters C5 and C6, horizontal and amacrine cluster C7, photoreceptors clusters C8 and C9, but little overlaps between the naïve RPCs and fate-committed neurons (Figure 4b, and data not shown), further confirming that C3 and C4 were in a transitional state linking naïve RPCs and differentiating neurons. Of note is that C10, which was composed of the ciliary margin cells with a unique gene signature, also expressed many of the genes enriched in RPC clusters C0-C2 (Figure 4a, Suppl Table 6), highlighting the close developmental relationship of the ciliary margin and the neural retina.

To further corroborate the relationships between the cells in these clusters, we also performed trajectory analysis using the SCANPY tool which is based on diffusion pseudotime (DPT) by measuring transitions between cells using diffusion-like random walks ^58^. As shown in Figure 4c and consistent with the conclusions from the heatmap with the cluster-enriched genes, three definitive trajectories representing photoreceptors, horizontal and amacrine cells, and RGCs were identified, which all originated from the transitional RPCs that were downstream of the naïve RPC clusters. Not surprisingly, being the first cell types to form, the RGC trajectory advanced the furthest (Figure 4c). All three trajectories also existed in the *Atoh7*-null cells (Figure 4d); whereas the other two trajectories were not affected, the RGC trajectory appeared to have progressed through C5 but stalled at C6, consistent with the fact that RGCs are specifically affected in the *Atoh7*-null retina.

### Characteristics of individual cell states during retinal development

To better understand the properties of the cell states/types represented by individual clusters, we further examined the biological function of the enriched genes in these clusters by GO analysis of each of these lists ^62^. For simplicity, we combined similar clusters, including C0 to C2 (naïve RPCs), C3 and C4 (transitional RPCs), C5 and C6 (RGCs), and C8 and C9 (photoreceptors) (Table 1). The top five GO biological processes enriched in naïve RPCs were cell cycle, cell division, mitotic nuclear division, nucleosome assembly, and chromosome segregation, confirming that they were indeed actively dividing RPCs at different phases of the cell cycle (Table 1). Two of the top five GO terms associated with C3 and C4 included cell cycle and cell division, further implying they were still RPCs. Interestingly, the other three top GO terms were all associated with RNA processing (Table 1), indicating that this process plays a critical role in these transitional RPCs. In contrast, GO terms enriched in RGCs, horizontal and amacrine cells, and photoreceptors were all related to the various aspect of neural development and function, further confirming that their identities were correctly assigned and that the enriched genes were involved in their formation.

**Table 1.** Enriched GO terms for individual cell states/types

| Term | Count | % | Fold Enrichment | P Value |
| --- | --- | --- | --- | --- |
| **Naïve RPCs (C0-C3)** |  |  |  |  |
| cell cycle | 64 | 18.66 | 5.84 | 2.63E-30 |
| cell division | 51 | 14.87 | 7.63 | 2.13E-29 |
| mitotic nuclear division | 42 | 12.24 | 8.49 | 6.07E-26 |
| nucleosome assembly | 22 | 6.41 | 11.84 | 1.20E-16 |
| chromosome segregation | 16 | 4.66 | 10.06 | 4.83E-11 |
| **Transitional RPCs (C3, C4)** |  |  |  |  |
| RNA splicing | 35 | 10.42 | 8.42 | 1.85E-21 |
| cell cycle | 51 | 15.18 | 4.81 | 2.52E-20 |
| mRNA processing | 37 | 11.01 | 6.66 | 3.06E-19 |
| cell division | 34 | 10.12 | 5.27 | 1.29E-14 |
| mRNA splicing, via spliceosome | 19 | 5.65 | 9.83 | 7.01E-13 |
| **RGCs (C5, C6)** |  |  |  |  |
| nervous system development | 63 | 7.67 | 4.07 | 3.24E-21 |
| axon guidance | 31 | 3.78 | 5.07 | 3.39E-13 |
| axonogenesis | 25 | 3.05 | 5.69 | 7.00E-12 |
| neuron projection development | 24 | 2.92 | 4.21 | 1.07E-08 |
| substantia nigra development | 13 | 1.58 | 8.34 | 2.08E-08 |
| **Photoreceptors (C8, C9)** |  |  |  |  |
| nervous system development | 30 | 8.17 | 4.35 | 7.13E-11 |
| axon guidance | 16 | 4.36 | 5.87 | 9.73E-08 |
| cell differentiation | 34 | 9.26 | 2.38 | 6.70E-06 |
| neuron migration | 12 | 3.27 | 5.29 | 1.71E-05 |
| multicellular organism development | 38 | 10.35 | 2.02 | 6.55E-05 |
| **Amacrine and Horizontal Cells (C7)** |  |  |  |  |
| visual perception | 13 | 3.80 | 5.99 | 1.82E-06 |
| photoreceptor cell maintenance | 7 | 2.05 | 10.73 | 4.23E-05 |
| synaptic vesicle exocytosis | 5 | 1.46 | 16.13 | 2.19E-04 |
| negative regulation of transcription from RNA polymerase II promoter | 26 | 7.60 | 2.19 | 3.79E-04 |
| positive regulation of transcription from RNA polymerase II promoter | 32 | 9.36 | 1.97 | 4.08E-04 |

The enriched gene lists also included many genes with unknown expression patterns and functions in the retina. To examine how faithful these enriched genes reflected their actual expression patterns in the developing retina, we chose genes from the enriched lists whose expression and function have not been well analyzed and compared their predicted expression patterns as presented by feature plots with that reported in the Eurexpress in situ hybridization database ^96^ (http://www.eurexpress.org/ee/). We found that the feature plots almost always correctly predicted the actual expression patterns, often with more details than in situ hybridization, as exemplified by 5 naïve RPC enriched genes and 10 RGC enriched genes whose expression and function in the retina have not been characterized (Suppl. Figures 2 and 3, Suppl. Table 6). Thus, the clustering data based on scRNA-seq analysis can serve as a very useful resource for identifying novel genes as markers or candidates for further functional analysis.

The scRNA-seq data also clarified contradicting results on two genes critical in retinal development. *Sox4* and *Sox11*, which encodes two transcription factors critical for RGC development ^97–99^, have been identified to be expressed mostly in RGCs, but there have been conflicting reports regarding whether they are also expressed in RPCs ^59,98–100^. In the *Atoh7*-null retina, since most RGCs are absent, it was indicated that *Sox4* and *Sox11* were downregulated ^59,98^. However, we did not detect significant changes in conventional RNA-seq analysis (Suppl. Table 2). These contradicting results were resolved by comparison of their expression in corresponding clusters; both genes were extensively expressed in all the clusters (Figure 5a, b), but were at lowest levels in naive RPCs (C0-2), began to increase in the transitional RPCs (C3, C4), and reached the highest levels in RGCs (C5, C6) and amacrine and horizontal precursors (C7) (Figure 5c). These patterns were further confirmed by in situ hybridization with RNAscope probes ^101^ (Figure 5d, e). In the *Atoh7*-null retina, the overall patterns of *Sox4* and *Sox11* remained and the levels were comparable in all clusters, with only moderate upregulation of *Sox4* in differentiated RGCs (C6, Figure 5c). Therefore, despite the loss of RGCs, the overall expression levels of these two genes did not change in the *Atoh7*-null retina as detected by regular RNA-seq using total RNA from whole retinas. Nevertheless, as further discussed later, the differential expression levels of *Sox4* and *Sox11* along the differentiation trajectories may be related to their functions in the retina, particularly in RGC genesis.

**Figure 5.**
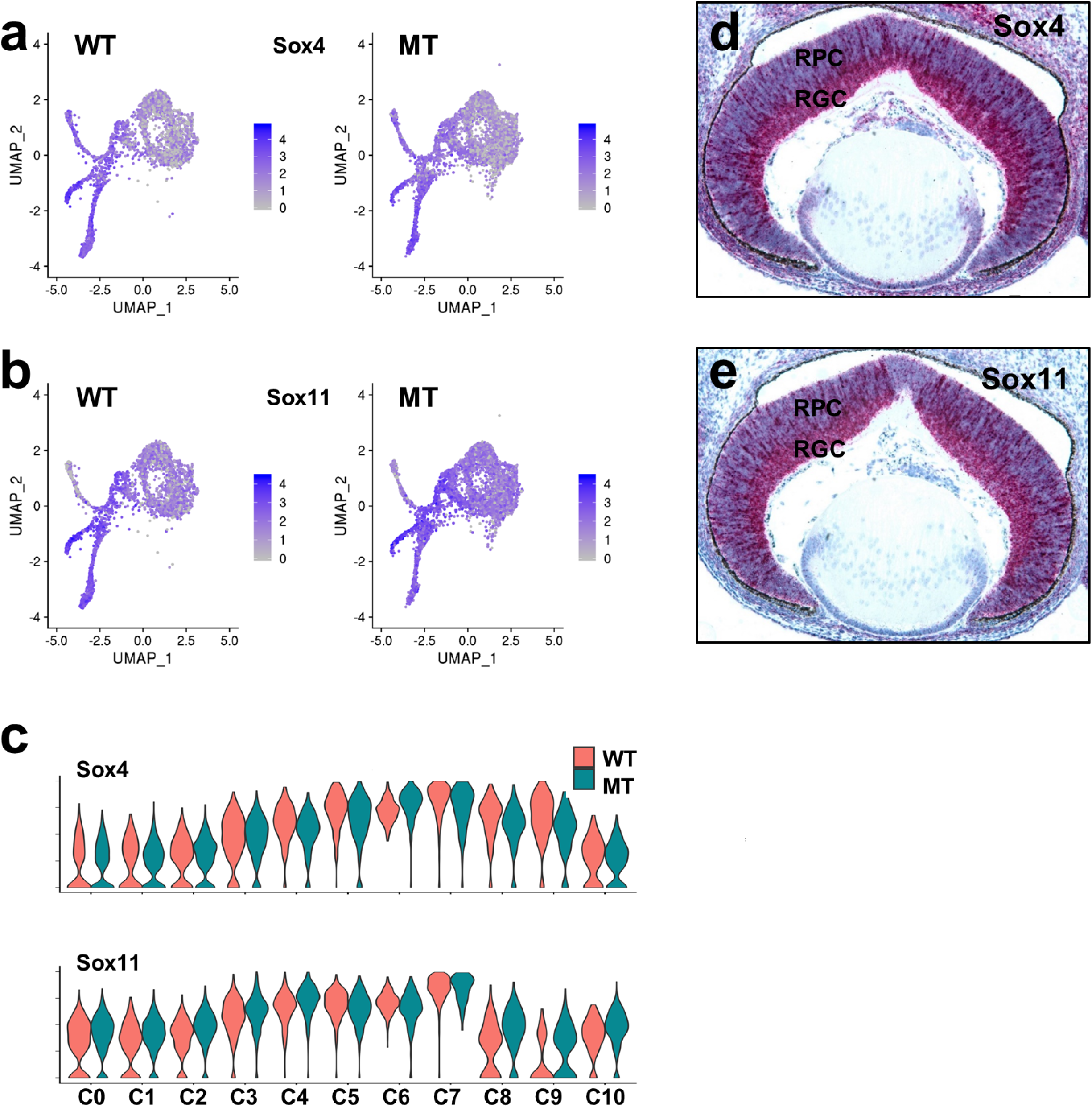
Spatial expression of *Sox4* and *Sox11*. **a**. Feature plot heatmaps indicate that both genes are expressed in all clusters but at much higher levels in differentiating cells, and that their expression levels are comparable in wild-type (WT) and *Atoh7*-null (MT) cells. **b**. Violin plots indicate that both genes are expressed in comparable levels in corresponding WT (red) and MT (blue) clusters. The heights of the plots represent expression levels and the widths represent relative proportions of cells expressing the gene at that level. **d**. **e**. In situ hybridization with RNAscope probes confirms that *Sox4* and *Sox11* are expressed in both RPCs and RGCs. Note that both genes are highly expressed in a subset of RPCs in the outside part of the retina, which presumably are the transitional RPCs.

### Atoh7 marks a transient state shared by all early differentiation cell fates

Cells in C3 and C4 appeared to represent a critical transitional stage linking naïve RPCs and differentiating neurons along individual lineage trajectories. They were considered RPCs since they still were in the cell cycle (Figure 3d, e), expressed many RPC marker genes (Figure 4a, b, Suppl. Table 6), and their fate was not committed (Figure 4c). Nevertheless, expression of many of the general RPC markers genes was significantly decreased in these cells (Figures 3c, 4a). Close examination indicated that these cells expressed many genes involved in specific cell lineages, including *Atoh7, Sox4, Sox11, Neurog2, Neurod1, Otx2, Onecut1, Foxn4, Ascl1, Olig2, Dlx1, Dlx2*, and *Bhlhe22* (Figure 6a and Suppl. Table 6). These genes have all been reported to be expressed in subsets of RPCs, and function in or mark specific lineages. For example, *Atoh7* and *Neurog2* are required for the RGC lineage ^31,32,102^, *Otx2* and *NeuroD1* function in the photoreceptor lineage ^23,95^, whereas *Olig2*-expressing cells give rise to cone and horizontal progenies ^19^. *Onecut1* and *Onecut2* function in essentially all the early retinal cell lineages ^26,29^. Other such genes included *Foxn4* for horizontal and amacrine cells, *Dlx1* and *Dlx2* for RGCs and amacrine cells, and *Bhlhe22* (also known as *bHLHb5*) for amacrine cells and bipolar cells ^21,63,103–105^.

**Figure 6.**
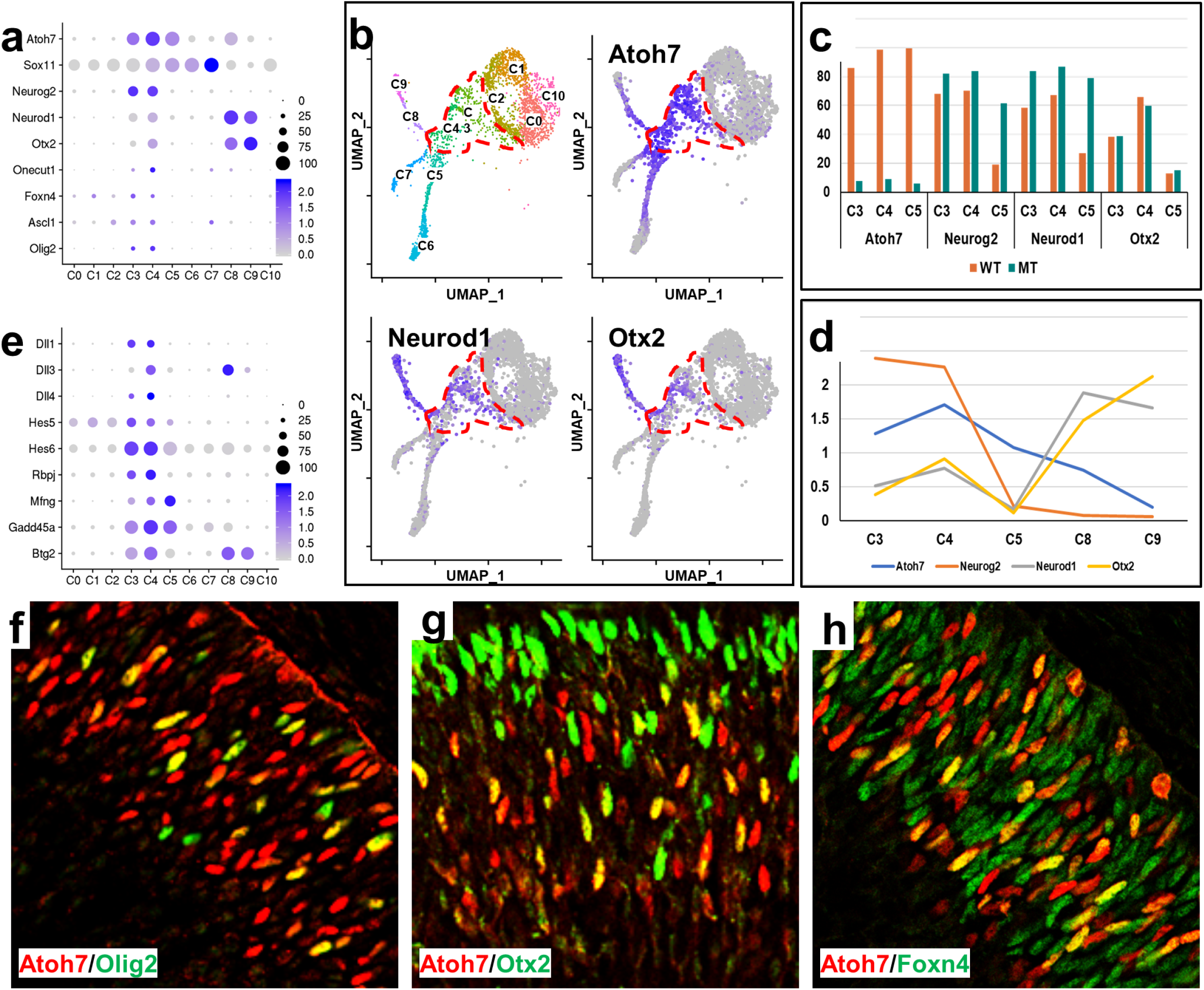
Gene expression signature of transitional RPCs (C3, C4). **a**. Genes regulating distinct lineages are expressed in a shared transitional state, namely transitional RPCs, as demonstrated by a dot plot. Often these genes continue to be expressed in the specific cell lineage they regulate, e.g. *Atoh7* and *Sox11* in RGCs (C5), and *Otx2* in photoreceptors (C8, C9). **b**. Feature heatmap showing the expression of *Atoh7*, *Neurod1*, and *Otx2* in the wild-type clusters. The dotted line demarcates the transitional RPCs (C3 and C4). Consistent with the dot plot, all three genes are expressed in transitional RPCs, but *Atoh7* is expressed in more transitional RPCs than *Neurod1* and *Otx2*. Whereas *Atoh7* trails into all three lineages, *Neurod1* and *Otx2* only continue to be expressed in photoreceptors. **c**. Percentage of cells expressing *Atoh7, Neurog2, Neurod1*, and *Otx2* in the transitional RPCs (C3, C4) and nascent RGCs (C5) in the wild-type (WT) and *Atoh7*-null (MT) retinas. **d**. Changes in activities of *Atoh7*, *Neurog2*, *NeuroD1*, and *Otx2* as cells progress into distinct lineages. For each gene, the gene activity in each cluster is derived by the expression level of that cluster divided by the mean of all clusters. **e**. Additional genes, including many encoding components of the Notch pathway, are enriched in the transitional RPCs (C3, C4). **f-h**. Immunofluorescence staining shows substantial co-expression of Atoh7 with Olig2 (**f**), Otx2 (**g**), and Foxn4 (**h**) in RPCs. Note the yellow cells are those expressing both markers in each panel. In the case of Otx2, which is expressed in both RPCs and photoreceptors (**g**), the co-expression only occurs in the RPCs.

The fact that these cells were clustered together and were enriched in these genes for different lineages indicated that they possessed shared properties. More interestingly, *Atoh7* was expressed in almost all cells in C3 (86%) and C4 (99%) and trailed into all three differentiating lineages (Figure 6a, b, c). Although this was consistent with previous findings that *Atoh7*-expressing cells are not fate-committed and can adopt all retinal fates ^31,32,46^, the high percentage of transitional RPCs expressing *Atoh7* was unexpected and likely significant. Moreover, several other factors such as *Neurog2*, *Neurod1*, and *Otx2* were also expressed in substantial portions of cells in C3 (68%, 58%, and 38% respectively) and C4 (70%, 67%, and 66% respectively) (Figure 6a, b, c). Noticeably, in the *Atoh7*-null retina, the proportions of *Neurog2*- and *Neurod1*- expressing transitional RPCs (C3 and C4) only increased slightly, and that of Otx2-expressing transitional RPCs did not change (Figure 6c). *Neurog2* and *Neurod1* were essentially turned off in wild-type nascent RGCs (C5), but remained highly expressed in corresponding *Atoh7*-null cells (Figure 6c).

As the trajectories of individual lineages progressed, the relative activities of these genes changed accordingly. In the nascent RGCs (C5), whereas *Atoh7* continued to be expressed at high levels, *Neurog2, Neurod1*, and *Otx2* were much reduced in expression (Figure 6a, b, d). On the contrary, in the photoreceptor lineage (C8 and C9), *Atoh7* and *Neurog2* levels dropped significantly, but *Neurod1* and *Otx2* increased markedly (Figure 6a, b, d). As mentioned above, *Sox4* and *Sox11*, had elevated expression in the transitional RPCs (C3 and C4) and continued to be expressed in all lineages (Figures, 5a-c, 6a). Nevertheless, other factors, such as *Olig2*, *Onecut1*, *Foxn4*, *Ascl1*, seemed to be expressed in considerably fewer transitional RPCs (Figure 6a). However, the overlaps between these genes could be more extensive as the percentage of gene expression in each cluster was likely underestimated due to sequence depth and expression levels. Nevertheless, the transitional RPCs likely remained heterogeneous.

Another prominent feature of the transitional RPCs is that many genes encoding components of the Notch pathway, including *Dll1*, *Dll3*, *Dll4*, *Notch1*, *Hes5*, *Hes6*, and *Mfng* were enriched, further emphasizing the critical roles this pathway plays in retinal development (Figure 6e, Suppl. Table 6). The expression of *Dll1*, *Dll3*, and *Dll4* was of particular interest; they all were only expressed highly in transitional RPCs (C3 and C4) and the differentiating clusters, but not much in the naïve RPC clusters. Although Hes5, one of the effector genes of the pathway, was enriched, *Hes1*, another downstream effector of the pathway, was significantly downregulated in C3 and C4, as compared to the naïve RPCs (Suppl. Table 6 and data not shown), indicating that these two genes were differentially regulated and likely had both shared and distinct functions ^74,106^. These findings suggested that when selected RPCs were poised for differentiation and began to express *Atoh7* and genes for other fates, they also elevated the levels of ligands of the Notch pathway, which in turn modulate the Notch activities in the naïve RPCs. Likely this is part of the mechanism by which the balance between proliferation and differentiation is achieved.

Many additional genes, e.g. *Gadd45a*, *Btg2*, *Penk*, *Srrm4*, and *Plk1, Sstr2, and Ccnb1*, were enriched in the transitional RPCs (Figure 6e, Suppl. Table 6), but their roles are mostly unknown. For example, *Gadd45a* and *Btg2*, two genes involved in cell cycle arrest, DNA repair and apoptosis ^107–110^, were highly enriched in transitional RPCs, but they diverge in the differentiating lineages (Figure 6e). *Gadd45a* continued to be expressed in nascent RGCs (C5), whereas *Btg2* maintained its expression in photoreceptors (C8, C9) (Figure 6e). We further confirmed by RNAscope in situ hybridization that they each indeed were expressed in a subset of RPCs at different developmental stages examined (E12.5, E14.5, and E17.5), with patterns very similar to that of *Atoh7* ^18,77,111^ (Suppl. Figure 4). Both genes are responsive to stress-induced growth arrest and inhibit the G1/S progression in the cell cycle. Although their roles in retinal development have not been well studied, they likely participate in establishing the transitional cell state in these cells, which are primed to exit the cell cycle and commit to distinct cell fates.

Our findings that *Atoh7* and genes for other cell lineages co-express in the transitional RPCs were consistent with previous co-immunofluorescence staining showing that Atoh7 overlaps substantially with multiple relevant factors such as Neurog2, Neurod1, Onecut1, and Onecut2 ^20,56,111^. Further, by co-immunofluorescence staining, we confirmed that large proportions of Olig2-, Otx2- and Foxn4-expressing RPCs (67.53±4.66%, 60.43±9.47 and 45.03±5.39 respectively, n=4) also expressed Atoh7 (Figure. 6f-h). The relatively low percentage of Foxn4 positive cells expressing Atoh7 likely was due to Foxn4 also being expressed in naïve RPCs (Figure 6a). These results indicated that all the early retinal cell fates go through a shared cell state which is characterized by downregulation of naïve RPC genes including those for cell cycle progression, upregulation of the Notch ligands, downregulation of the Notch pathway, and upregulation of neurogenic genes for various retinal fates. Of note is that, although *Atoh7* was expressed in essentially all transitional RPCs, deletion of *Atoh7* did not affect the formation of this cell state (Figures 3b, 6c), indicating that Atoh7 is not required for the establishment of this critical cell state in retinal development.

### Cell cluster-specific changes in gene expression in the *Atoh7*-null retina

Corresponding E13.5 wild-type and *Atoh7*-null clusters almost completely overlapped in the 2D projection of the UMAP analysis (Figure 3a, b). Most clusters, including the transitional RPCs (C3 and C4), contained comparable proportions of cells in the wild-type and mutant retinas (Figure 7a). However, several clusters demonstrated marked changes. There was an about two-fold increase in the proportion of mutant C0 cells. Since most C0 cells were in G1/S of the cell cycle (Figure 2c, d), this may reflect the reduced proliferation of the naive RPCs caused by reduced Shh signaling and the G1/S cyclin (Cyclin D1) levels (Figure 2) ^22,31,66,68,112^, but the other pathways including the Notch pathway may also be involved. The cell number in mutant C7 reduced almost by half, which is consistent with previous reports suggesting that Atoh7 plays a role in the genesis of horizontal cells ^31,59^. There was also a noticeable drop in the number of nascent photoreceptor cells (C8), but no change in the more differentiated photoreceptors (C9). The significance of this observation is not known. The mutant cluster with the most significant change was C6, which were differentiating RGCs, with an ~5 fold reduction as compared to the wild-type cluster (Figure 7a). However, the nascent RGC cluster (C5) and the transitional RPC clusters (C3 and C4) did not show obvious changes in cell numbers. As discussed earlier, deletion of *Atoh7* did not substantially affect the overall cell cycle status of each cluster except for C0 as noted above, the relationships of the different clusters, or the overall trajectories of the distinct lineages (Figure 3a-e, Figure 4c, d).

**Figure 7.**
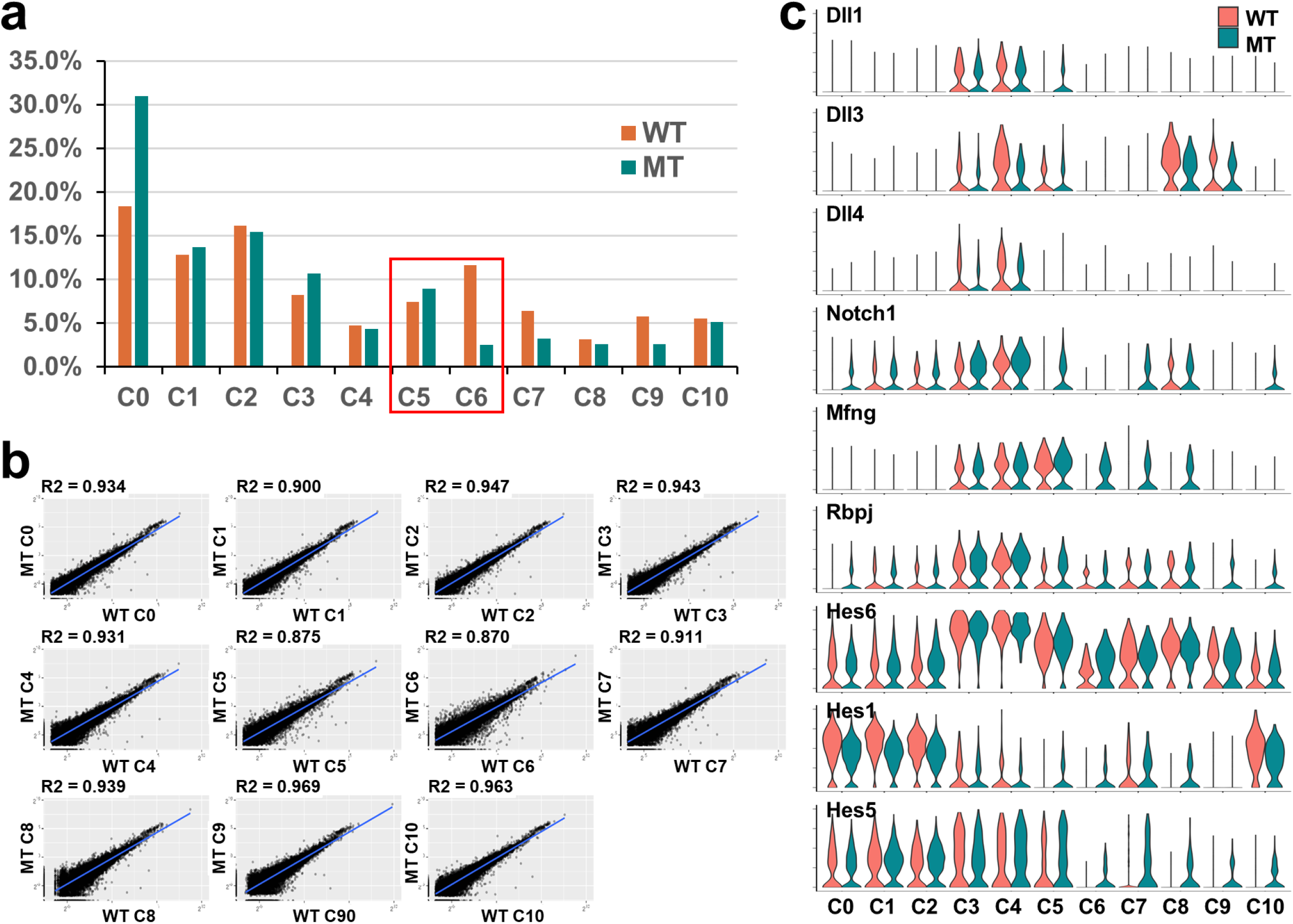
Comparison of wild-type and *Atoh7*-null clusters. **a**. Proportions of cells in each wild-type (WT, red) and *Atoh7*-null (MT, blue) clusters. Nascent RGCs (C5) and differentiated RGCs (C6) are highlighted by a red box. Note there is no major change in the proportion of cells in C5, but a marked reduction in C6. **b**. Scatter plots comparing gene expression in corresponding pairs of WT and MT clusters. The correlation coefficients (R^2^) are shown for each pair. The C5 and C6 pairs have the lowest R^2^ values, indicating the most changes in gene expression. **c**. Violin plots showing cluster-specific changes in expression of the Notch pathway genes. Note the distinct expression patterns of individual genes and differential expression changes in the MT clusters (see text for details).

The same corresponding clusters observed in the *Atoh7*-null retina provided us the opportunity to probe the cell state/type-specific gene expression changes caused by deletion of *Atoh7*. Global gene expression between corresponding wild-type and *Atoh7*-null clusters were highly similar as revealed by the scatter plots; the correlation coefficients (R^2^) ranged from 0.870 to 0.969 (Figure 7b). The high R^2^ values indicated that gene expression levels detected by scRNA-seq were highly robust and reproducible. They were also consistent with the knowledge that Atoh7 functions highly specifically in the RGC lineage. Accordingly, the two RGC clusters (C5 and C6) had the lowest R^2^ values (0.875 and 0.870 respectively). By comparing corresponding wild-type and *Atoh7*-null clusters, we identified a total of 1829 DEGs (Suppl. Tables 8-18). Comparison with the DEG list from conventional RNA-seq revealed an overlap of only 290 genes (Suppl. Tables 2, 8-18). Further examination of the none-overlapped genes indicated scRNA-seq could not effectively detect DEGs with relatively low expression levels. For example, *Shh* was readily detected as a DEG by regular RNA-seq, but not by scRNA-seq, although *Gli1*, the downstream target gene of the Shh pathway, was detected by both methods (Figure 2a, Suppl. Tables 2, 8-10, 13, 14). This was likely due to the relatively low sequencing coverage of the transcriptome in scRNA-seq. Conventional RNA-seq, on the other hand, was inefficient in detecting DEGs that were expressed in multiple clusters but the change only occurred in selected clusters (see below). These results indicated that each method had limitations in identifying DEGs, particularly in tissues with complex cellular compositions, and that the two methods were complementary in providing a more complete picture of changes in gene expression.

Nevertheless, genes identified by comparing the corresponding wild-type and mutant clusters provided further insights into the function of Atoh7. Our focus will be on the DEGs in naïve RPCs (Suppl. Tables 8-10), transitional RPCs (Suppl. Tables 11, 12), and RGCs (Suppl. Tables 13, 14), since they were directly related to RGC development, whereas only small numbers of DEGs were detected for other clusters (Suppl. Tables 15-18). Consistently, GO analysis of downregulated DEGs in naïve RPCs (C0-2) by DAVID revealed the top enriched biological process GO terms included protein folding, mRNA processing/RNA splicing, and cell division/cell cycle (Suppl. Table 8-10, 19). These biological processes are all required for active proliferation, which is a property of naïve RPCs. Example DEGs directly involved proliferation included *Ccnd1, Lig1, Mcm3*, and *Mcm7*. On the other hand, one of the prominent features of the biological processes associated with the upregulated DEGs is gene regulation associated with neural development (Suppl. Table 19). These results suggested that there was a shift in the properties of the naïve RPCs from proliferation to differentiation. Since *Atoh7* is not expressed in these cells (Figure 3c, Figure 6b), this shift likely was caused by non-cell autonomous mechanisms, highlighting the interaction between RGCs and RPCs. DEGs in the transitional RPCs (C3 and C4) likely reflected its direct function. Downregulated genes included those implicated in the RGC lineage such as *Dlx2* and *Eya2*, those involved in mRNA processing, and the Delta-like ligand genes (see below), but not cell cycle genes (Suppl. Tables 11, 12, 19). On the other hand, upregulated DEGs included those encoding a large number of transcription factors, many of which, such as *Foxn4, Neurod1*, and *Onecut2*, are involved in non-RGC lineages.

As expected, the largest number of DEGs were found between the wild-type and mutant RGC clusters (C5 and C6); 450 DEGs were found in C5, 618 DEGs were found in C6, and collectively a total of 861 DEGs were identified in these two clusters (Suppl. Tables 13, 14). These genes are directly relevant to the establishment and maintenance of the RGC identity; among them included *Pou4f2* and *Isl1* and many other previously identified genes encoding regulatory, functional, and structural proteins critical for RGC differentiation, which was further confirmed by GO analysis (Suppl. Table 19) ^49,50,59,60^. As further discussed later, the upregulated DEGs in the mutant RGCs featured a large set of genes normally expressed in RPCs and/or other cell types including those of the Notch pathway (Suppl. Tables 13, 14). These upregulated DEGs indicated the RGC lineage, although still formed in the *Atoh7*-null retina, was immature and likely had mixed identities.

One interesting observation from the cluster-specific comparisons of the scRNA-seq data was that the Notch pathway was affected in a complex fashion in the *Atoh7*-null retina (Figure 7c, Suppl. Tables 8-17). In the naïve RPCs (C0-C2), *Hes1*, but not *Hes5*, was significantly down-regulated (Figure 7c, Suppl. Table 8-10), indicating the downregulation of the Notch pathway. Since *Hes1* is a convergent signaling node ^66,68,73^, its downregulation likely resulted from the combined dysregulation of multiple pathways, including the Shh pathway, the Vegf pathway, and the Notch pathway its self. Consistently, in the transitional RPCs (C3 and C4), all three delta-like ligand genes, *Dll1, Dll3*, and *Dll4*, were markedly reduced, but other genes enriched in these cells, including *Notch1, Mfng*, and *Rbpj*, and *Hes6* did not change (Figure 7c, Suppl. Tables 11, 12). On the other hand, multiple Notch pathway genes, including *Dll1*, *Dll3, Notch1, Mfng*, *Rbpj*, *Hes1*, and *Hes5*, were upregulated in RGCs (C5 and/or C6) (Figure 7c, Suppl. Tables 15, 16). Thus, Atoh7, which is expressed in the transitional RPCs, may influence the Notch pathway in multiple cell types, likely through regulating the Delta-like ligand genes directly and the other signaling pathways indirectly. As suggested above, the continued expression of Notch components in the RGCs may reflect their immaturity and stalled differentiation.

It is worth noting that the functions of many of the DEGs, both down- and upregulated, are unknown. Such examples included genes encoding members of the chaperonin containing TCP1 complex (*Cct3, Cct4, Cct5, Cct6A, Cct7, Cct8*), proteins involved in mRNA processing, and many transcription factors (e.g. *Insm1, Id1, Id2, and Id3*) (Suppl Tables 8-17). Further investigations are needed to understand their roles in the developing retina.

### The RGC lineage forms and advances substantially in the absence of Atoh7

The two clusters representing the RGC lineage (C5 and C6) were still present in the *Atoh7*-null retina (Figures 3b, Figure 7a). Although a significant reduction in cell number was found in more differentiated RGCs (C6) as expected, no major change in the proportion of nascent RGCs (C5) was observed in the *Atoh7*-null retina (Figure 7a). Thus, the RGC lineage was still formed initially, but its developmental trajectory stalked prematurely (Figure 4c, d, Figure 7a). These findings indicated that, contrary to previous conclusions ^25,30^, the RGC lineage still formed and advanced considerably in the absence of Atoh7, although most cells eventually failed to differentiate into fully functional RGCs.

To better understand the underlying genetic mechanisms for the defective RGC development in the *Atoh7*-null retina, we further examined the DEGs in C5 and C6 by comparing them (861 genes, Suppl. Table 13, 14) with the enriched genes in C5 and C6 (821 genes, Suppl. Table 6), which were mostly RGC-specific genes (Figure 8a). Although a significant portion of the C5/C6 enriched genes (386 genes) were downregulated as expected, many of them did not change in expression (405 genes). A small number of C5/C6-enriched genes (34 genes) were upregulated. On the other hand, many downregulated DEGs (198 genes) and most upregulated DEGs (246 genes) were not enriched in C5/C6. These findings suggested these genes were regulated in different modes in the RGC lineage.

**Figure 8.**
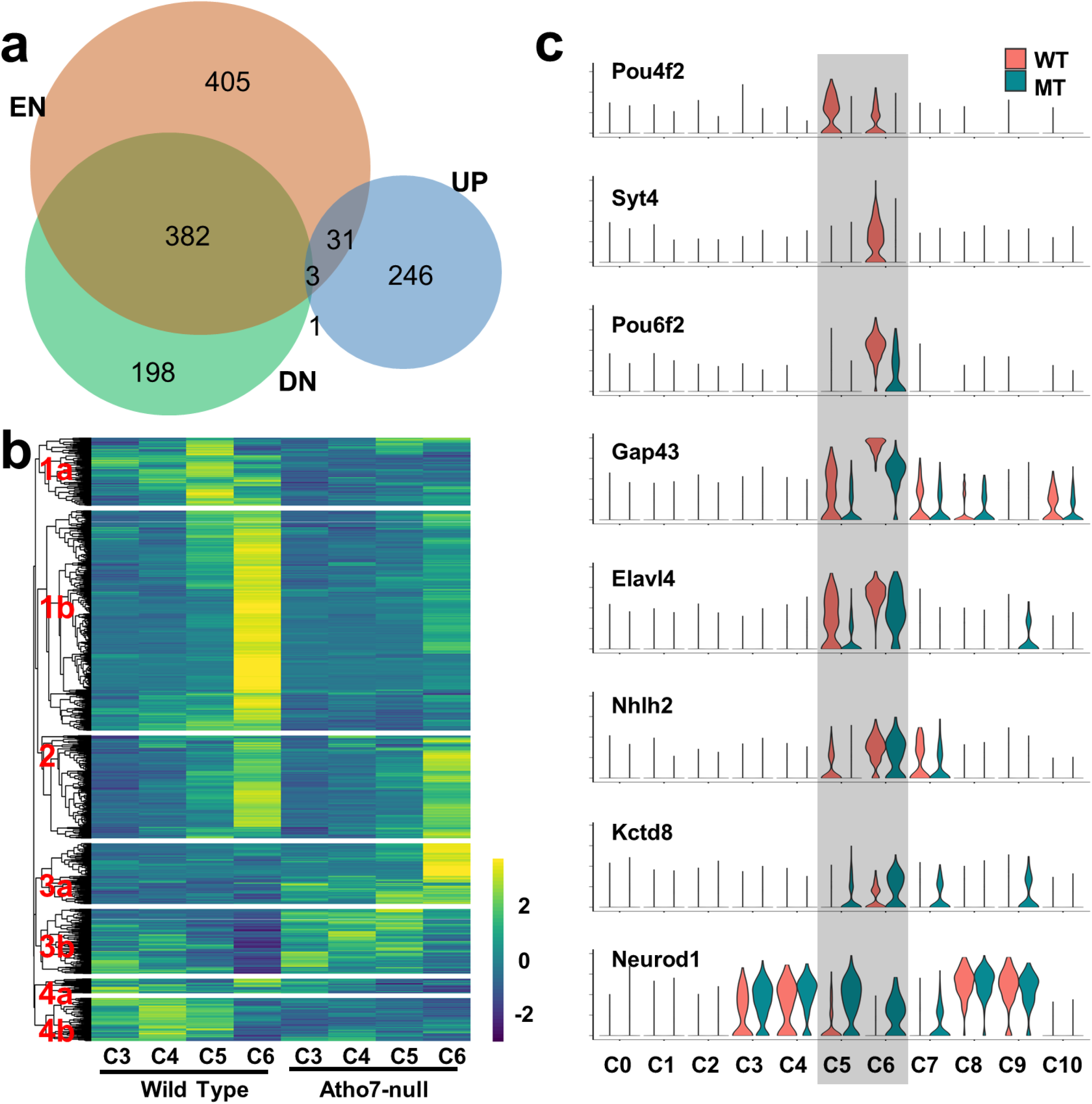
Gene expression underlying the RGC lineage in the *Atoh7*-null retina. **a**. Overlaps of down- and upregulated DEGs (DN and UP respectively) in *Atoh7*-null RGCs and RGC-enriched genes (EN) as presented by a Venn diagram. **b**. Euclidean distance clustering demonstrating seven different modes of changes in gene expression in *Atoh7*-null retinal RGCs (C5 andC6, see main text for details). For comparison, expression in C3 and C4 are also presented. **c**. Example genes with different modes of expression as demonstrated by violin plots, showing their expression across all clusters of wild-type (WT, red) and *Atoh7*-null (MT, blue) cells. *Pou4f2* is from group 1a in B, *Syt4, Pou6f2, Gap43*, and *Elavl4* are from group 1b, *Nhlh2* is from group 2, *Kctd8* is from group 3a, and *Neurod1* is from group 3b.

To further confirm these findings, we performed unsupervised clustering of all the 1266 genes included in the C5/C6 DEG list and enriched gene list across the clusters of wild-type and *Atoh7*-null cells. This led to 7 groups of genes with distinct expression dynamics across the clusters, including RGC-enriched and downregulated (Group 1a, b); RGC-enriched but not significantly changed (Group 2); none RGC-enriched but upregulated (Group 3a, b); and none RGC-enriched but downregulated (Group 4a, b) (Figure 8b, only C3 to C6 are shown, and Suppl. Table 20). Example genes representing some of these distinct expression modes across different cell types/states were more clearly demonstrated by violin plots (Figure 8c). From these analyses, it was apparent that, as expected, a large proportion of RGC-specific genes, including *Pou4f2*, *Isl1*, *Syt4*, *Pou6f2*, *Gap43*, *Elavl4*, were downregulated in the *Atoh7*-null retina (Figure 8). Among them, some genes such as *Pou4f2* and *Syt4* exhibited little expression in the mutant, but other genes, such as *Isl1* (data not shown), *Pou6f2 Gap43*, *Elavle4*, were still expressed at variable but substantial levels (Figure 8b, c). In addition, a substantial number of RGC genes such as *Nhlh2* remained expressed at similar levels as in the wild-type retina (Figure 8b, c). The remaining RGC-specific gene expression in the *Atoh7*-null retina, albeit often at lower levels, likely underlay the presence of nascent and differentiating RGCs.

Nevertheless, these RGCs not only failed to express a large number of genes either completely or at sufficient levels, but also aberrantly overexpressed many genes not enriched in RGCs (Groups 3a, b in Figure 8b, Figure 8c). Some of these genes, e.g. *Kctd8* (Figure 8c), were expressed at low levels in all wild-type clusters but were significantly upregulated specifically in RGCs (Group 3a). Other genes expressed in naïve and transitional RPCs but not in RGCs in the wild-type retina remained expressed in the mutant RGCs (Group 3b). As mentioned earlier, among these genes included the Notch pathway genes, *Neurod1*, and *Neurog2*. *Neurod1*, which is normally expressed in the transitional RPCs (C3 and C4) and photoreceptors (C8, and C9), but not in RGCs (C5 and C6) (Figure 6c,d, Figure 8c) became highly overexpressed in the *Atoh7*-null RGCs (Figure 8c).

The collective dysregulation of genes in *Atoh7*-null cells likely led to the truncated RGC trajectory. These cells progressed to an RGC-like state by expressing many of the RGC genes, often at reduced levels, but also overexpressed many genes abnormally. Because of the aberrant gene expression, they failed to fully differentiate into more mature RGCs and many of them, if not all, likely died ^32^. Noticeably, in the *Atoh7*-null retina, we did not find increased expression of genes directly involved in apoptosis in either the regular RNA-seq or scRNA-seq. This likely was due to the relatively small number of dying cells at any given time and that very few dying cells, if any, contributed to the single cell libraries.

### Effects on retinal development at E17.5 by *Atoh7* deletion

To further investigate what occurred to the RGCs in the *Atoh7*-null retina as development proceeded, we also compared expression profiles of the wild-type and *Atoh7*-null retinas at E17.5, a time when RGC production significantly decreased ^4,18^. Since the proportions of cells in RGC lineage become increasingly smaller as development advances ^4,18^, we took advantage of two knockin-mouse lines, *Atoh7^zsGreenCreERT2^* and *Pou4f2^FlagtdTomato^*, which label Atoh7- and Pou4f2-expressing cells respectively, and enriched these cells by FACS ^52^. Due to the stability of the zsGreen protein, zsGreen cells also included progenies of Atoh7-expressing cells, including RGCs, horizontal cells, and photoreceptors. We enriched Atoh7-expressing cells from both heterozygous (*Atoh7^zsGreenCreERT2/+^*, WT) and null retinas (*Atoh7^zsGreenCreERT2/lacZ^*, MT), as well as RGCs from the *Pou4f2^FlagtdTomato/+^* (WT) retina. We used low gating thresholds to enrich the relevant cell populations, but not to exclude other cell types, which allowed us to profile all the cell populations at E17.5 (see below). For simplicity of our analysis, zsGreen cells from the retina *Atoh7^zsGreenCreERT2^* and RGCs from the *Pou4f2^FlagtdTomato^* retina were grouped together (28,283 cells) as they are phenotypically wild-type, and compared with *Atoh7*-null cells (17,175 cells). As done with E13.5 cells, UMAP projection clustering and marker analysis allowed us to identify cell groups with similar cell identifies, including naïve RPCs, transitional RPCs, horizontal and amacrine cells, and photoreceptors for both wild-type and *Atoh7*-null cells, although the exact numbers of clusters differed and the identity of one cluster could not be ascertained (Figure 9, Suppl. Figure 5). Similar relationships from naïve RPCs to transitional RPCs, and then to fate-committed retinal cell types, were observed with both wild-type and *Atoh7*-null cells (Figure 9a). Importantly, transitional RPCs continued to express genes for multiple cell types with high overlaps, which included *Atoh7, Neurog2, NeuorD1, Otx2*, *Foxn4*, and *Olig2* (Figure 9b and data not shown). These genes, except *Atoh7*, were also expressed in the *Atoh7*-null transitional RPCs (Figure 9b and data not shown). The other genes expressed in the E13.5 transitional RPCs, including *Gadd45a* and *Btg2*, also continued to mark these cells at E17.5 (Suppl. Figure 5), indicating the general properties of these cells remained, although the cell types they produced had shifted. The presence of the RGC cluster in the E17.5 *Atoh7*-null retina suggested that some of the mutant RGCs persisted for some time. We further validated this by immunofluorescence staining for two RGC markers, Nefm and Uchl1, which showed that about 10-15% RGCs remained in the *Atoh7*-null retina at E17.5 (Suppl. Figure 6).

**Figure 9.**
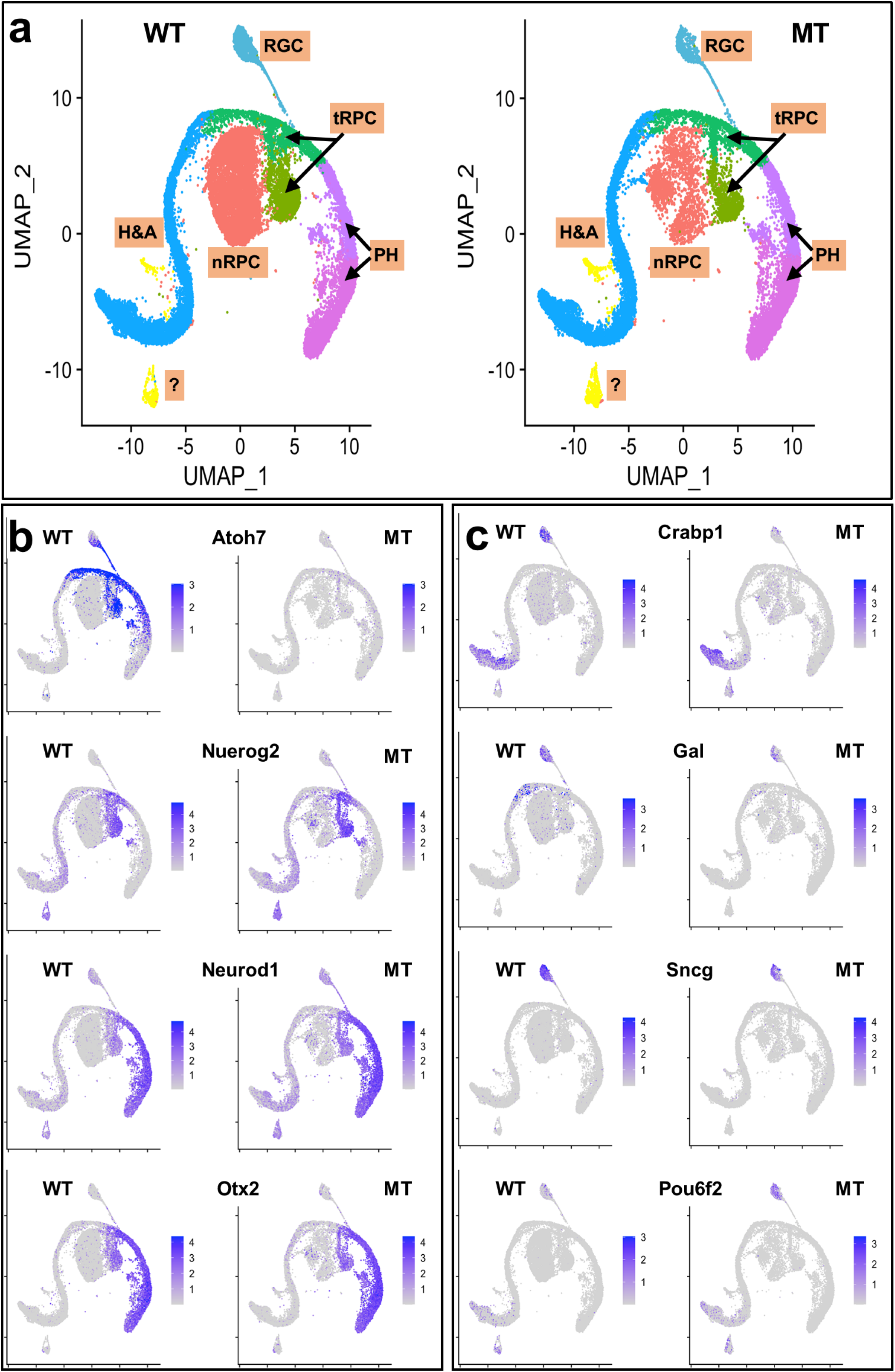
Single cell RNA-seq analysis of E17.5 wild-type (WT) and *Atoh7*-null (MT) cells. **a**. Identifies of UMAP clusters of WT and MT retinal cells, including naïve RPCs (nRPC), transitional RPCs (tRPC), RGCs, horizontal and amacrine cells (H&A), photoreceptors (PH). Note there are two transitional RPC clusters, two photoreceptor clusters, and a cluster (?) whose identity needs to be further confirmed. **b**. Feature heatmaps showing genes expressed in transitional RPCs, including *Atoh7*, *Neurog2*, *Neurod1*, and *Otx2* in WT and MT retinas. **c**. Feature heatmaps of a set of RGC genes in WT and MT retinas, including *Crabp1, Gal, Sncg*, and *Pou6f2. Crabp1, Gal*, and *Sncg* are down-regulated, *whereas Pou6f2* is up-regulated, in the MT RGCs.

Comparison of the same clusters between wild-type and *Atoh7*-null retinas revealed the changes in E17.5 naïve RPCs largely followed the trend at E13.5 with many of the same genes affected in both stages (Suppl. Tables 21). For example, genes involved in cell cycle progression, including *Ccnd1, Mcm7, Mcm3, and Hes1*, continued to be downregulated, likely due to RGC loss and disrupted signaling from them. In the transitional RPCs, fewer genes were affected at smaller fold changes, likely reflecting the reduced activity of *Atoh7* and RGC production at this stage ^4,18,77^, but many genes, such as *Dll3* and *Neurod1*, continued to be affected the same way as at E13.5 (Suppl. Tables 22, 23). Consistently and unlike at E13.5, both wild-type and *Atoh7*-null RGC clusters were only tenuously connected to the transitional RPC clusters at E17.5 (Figure 9a). In contrast to E13.5, there were only a small number of DEGs identified in the mutant RGCs at E17.5, and the fold changes tended to be smaller (Suppl. Table 24). However, these DEGs indicated that the remaining RGCs still were not normal. Whereas many RGC genes such as *Pou4f2, Elavl4, Nefm, Nefl* were expressed at the wild-type levels, other genes, e.g., *Crabp1, Gal, Sncg, Gap43, Ebf1, Ebf2, Ebf3*, Klf7, *Irx3, Irx5*, and *Isl1*, were downregulated. Intriguingly, *Pou6f2*, which was downregulated at E13.5 (fold change −2.9), was highly upregulated in the E17.5 *Atoh7*-null retina (fold change 2.5) (Figure 9c, Suppl Figure 6, Suppl. Tables 14, 24). Although the significance and mechanisms of the differential responses of the RGC genes in the *Atoh7*-null retina are unknown, they likely contributed to the eventual loss of almost all RGCs ^25,30^.

## Discussion

In this study, we first used regular RNA-seq to investigate the global transcriptomic changes in three mutant retinas, *Atoh7*-, *Pou4f2*-, and *Isl1*-null, during early development. The RNA-seq data provide a comprehensive list of genes expressed in the early developing retina (Suppl. Table 1). All genes known to function at this stage are on the list, and the gene list can be further mined for novel key regulators. Since all three mutants are defective in RGC development, this analysis provides a more complete picture and expands our knowledge of the function and hierarchical relationships of the three transcription factors in this lineage. The overlaps of downstream genes activated by the three factors confirm that Atoh7 acts upstream, whereas Pou4f2 and Isl1 are dependent on Atoh7 but only represent a part of the downstream events. Multiple signaling pathways are downstream of the three factors and functions through complex feedback mechanisms to coordinate proliferation and differentiation. Many RGC-specific genes were only dependent on Atoh7, but not on Pou4f2 or Isl1, indicating other factors parallel to Pou4f2 and Isl1 are at work in the RGC lineage. On the other hand, the lists of upregulated genes, which are normally repressed by the three transcription factors, demonstrated that the three factors exert their repressive roles largely independently at two levels: Atoh7 in RPCs whereas Pou4f2 and Isl1 in RGCs, although significant crosstalk between the two levels exists.

We then performed scRNA-seq on E13.5 and E17.5 wild-type and *Atoh7*-null retinal cells. The analysis not only correctly identified known cell states/types present at the stage, but also identified enriched genes for each cluster. The cluster-specific expression provided precise expression information not available before, demonstrating the power of this technology. Specifically, genes enriched in different clusters from our scRNA-seq data define specific states along the developmental trajectories and provide highly accurate information on their cell state/type-specific expression patterns (Figure 10). These results not only validate several recent reports of scRNA-seq analysis of both mouse and human retinas showing the presence of distinct lineage trajectories and a shared transitional but plastic (multipotent) cell state (transitional RPCs) by all the early trajectories ^13,38,39,44^, but also significantly extend those findings by revealing that *Aoth7* is expressed in all transitional RPCs and highly overlaps with genes involved in lineages other than RGCs. In agreement with our results, a similar finding that *Atoh7* marks transitional RPCs has also been reported with human embryonic retinal cells ^44^. Our study provides further insights into the nature of RPC competence for different retinal cell fates and the likely mechanism by which these fates are committed. Previous studies indicated that subsets of RPCs marked by specific genes exist and that these subsets are required for or biased toward particular cell fates ^17,19,31,32^. However, the relationship between these RPC subpopulations and its relevance to the competence of RPCs for different retinal fates have not been known. Our current study establishes that these populations highly overlap and can be considered as a shared cell state of all early retinal cell types (Figure 10). This state is characterized by co-expression of genes essential for individual retinal cell types, such as *Atoh7* and *Neurog2* for RGCs, *Otx2* and *Neurod1* for photoreceptors, and *Foxn4* and *Onecut1/2* for amacrine and horizontal cells. The commonality of these factors is that they function before fate commitment but promote RPCs toward individual lineages. The transitional RPCs, still dividing but likely in the last cell cycle(s) ^18,31,32,46,111,112^, are also characterized by significantly reduced expression of the naïve RPC markers and proliferation genes, increased expression of ligands for the Notch pathway, and decrease in the Notch pathway activities. These aspects of transition are likely coordinated, although the underlying mechanisms are not known. The Notch pathway is essential for RPC proliferation but inhibits differentiation ^5,106,113,114,114–118^. Promotion of proliferation by Notch may be achieved through interaction with some of the naïve RPC genes such as *Sox2, Lhx2*, and *Pax6* ^76,86,119,120^. Consistently, retinal cell differentiation requires the downregulation of the Notch pathway ^5,115,121,122^. Thus, downregulation of the Notch pathway likely is a key step for establishing this transitional state, and upregulation of the Notch ligands and other components may serve as a mechanism to balance proliferation and differentiation by lateral inhibition ^123–125^. This downregulation is likely mediated in part by transcription factors like Atoh7, Ascl1, and Foxn4 ^121,126,127^. Our results indicate that Atoh7 influences the Notch pathway in a complex fashion, both directly and indirectly, in different cell states/types. Additional genes, such as *Gadd45a, Btg2, Penk, Srrm4, Plk1, Sstr2, and Ccnb1*, were found highly expressed in this transitional state; they likely also play key roles in the transition from naïve RPCs to transitional RPCs.

**Figure 10.**
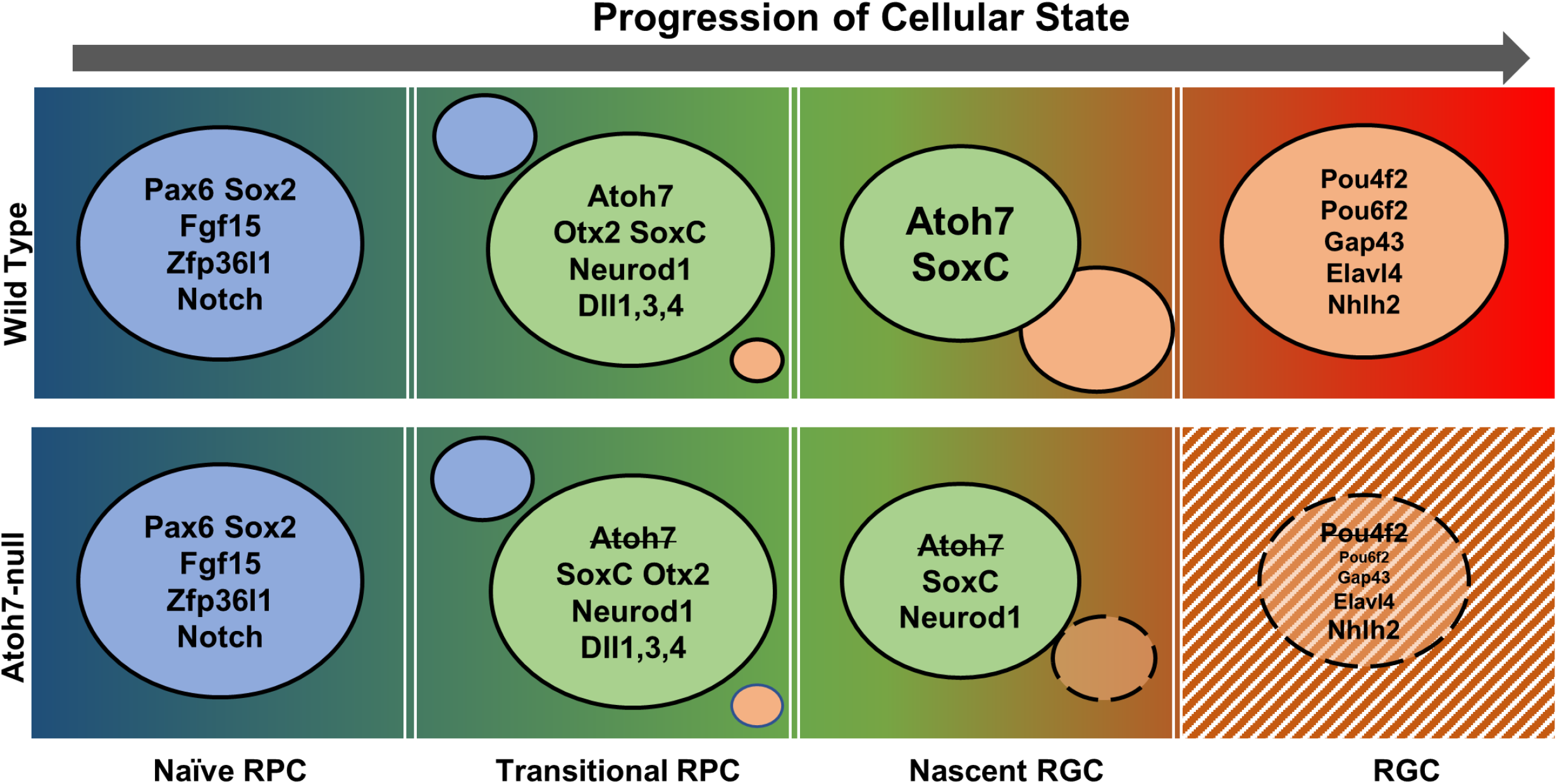
A model explaining shifts of cell states along the RGC trajectory in the wild-type and *Atoh7*-null retina. The RGC trajectory follows several four cell states, including naïve RPCs, transitional RPCs, nascent RGCs, and differentiated RGCs. The direction of the trajectory is indicated by the arrow and the progression of the trajectory is indicated by a color gradient. Each state is determined by a group of genes and example genes are given in colored circles. The transition from one state to the next is dictated by downregulation of genes representing that state and upregulation of genes for the next state as indicated by the sizes of the colored circles. In the transitional RPCs, Atoh7, likely in combination with the SoxC factors, competes with other regulators to drive them to the RGC fate. In the *Atoh7*-null retina, the establishment of the transitional RPC state is not affected, and nascent RGCs still form through expression of some, but not all, RGC genes (represented by a smaller circle with broken lines). The mutant nascent RGCs fail to reach the full RGC state and many eventually die by apoptosis (indicated by a striped background).

Since all early cell types arise from these transitional RPCs, as suggested by our trajectory analysis, the long-postulated RPC competence for retinal cell fates may be determined and defined by the genes expressed in them. For example, their competence for the RGC fate is dictated at least in part by *Atoh7*, whose expression coincides with RGC production ^18,77,111^. At later stages, when *Atoh7* is not expressed, RPCs lose their competence for the RGC fate. Consistent with the idea, deletion of *Atoh7* does not affect the establishment of this transitional state or the competences for other cell types. Since key regulators of different fates are co-expressed in these transitional RPCs, an outstanding question that arises is how the eventual outcome, i.e. adopting one particular fate versus another, is achieved. The mechanisms by which Atoh7 promotes the RGC fate may serve as a point of discussion regarding how transitional RPCs take on a specific developmental trajectory. In agreement with previous findings that Atoh7 is essential but not sufficient for the RGC lineage, we observed that *Atoh7* is expressed in all transitional RPCs and its expression trails into all three lineages being generated at E13.5. Since Atoh7 is expressed in all transitional RPCs and thus significantly overlap with competent factors for other fates (e.g. Neurod1 and Otx2 for photoreceptors), these factors likely compete with each other to steer the transitional RPCs toward different directions (Figure 10). The competition may occur at transcription levels through cross-repression as evidenced by the upregulation in the *Atoh7*-null retina of *Neurod1* and *Bhlhe22*, which are required for photoreceptors and amacrine cells respectively and by the distinct expression dynamics along different trajectories. However, this may not be the only or even the dominant mechanism, as other genes such as *Otx2* expressed in the transitional RPCs are not affected by the loss of Atoh7. Thus, Atoh7 and the other competence factors may also compete with each other stochastically by activating distinct sets of downstream genes essential for the respective fates, e.g. *Pou4f2* and *Isl1* for the RGCs, *Ptf1a* for horizontal and amacrine cells, and *Crx* for photoreceptors. This idea is consistent with previous observations that RPCs generated different retinal cell types in a stochastic fashion ^7,8^. Nevertheless, the eventual outcome is likely determined genetically; the proportion of different cell types produced at any given time is likely dictated by the presence of the competence factors and their relative activities. This idea is further supported by the finding that *Atoh7* gene dosage affects the number of RGCs produced and that overexpression of Atoh7 produces more RGCs ^128–130^. Activities of these transcription factors may also be modified posttranslationally ^131–134^. On the other hand, the transitional RPCs are still heterogeneous, as indicated by the uneven expression of many genes in these cells. This has also been demonstrated by lineage-tracing experiments; although *Olig2*, *Neurog2*, and *Ascl1* are all expressed in transitional RPCs, cells expressing these genes are biased in producing specific retinal progenies ^17,19^. The heterogeneity of transitional RPCs may reflect their different degree of progression toward different developmental trajectories.

Our scRNA-seq analysis on the *Atoh7*-null retina leads to significant insights into RGC development by identifying specific changes in gene expression through both direct and indirect mechanisms. Of particular significance was our observation that in the absence of Atoh7, the RGC trajectory still progressed considerably, but stalked prematurely. This may have been observed previously but not fully appreciated; many of the mutant Atoh7-expressing cells, marked by knock-in lacZ or Cre-activated reporter markers, still migrate to the inner side where RGCs normally reside ^25,30,31,46^, but many of them likely die by apoptosis ^32^, although some of these cells may redirect and adopt other fates. However, the status of these cells has not been well characterized. Our results indicate that these cells are on the RGC trajectory, and many, but not all, RGC-specific genes are activated in them, although often not to the wild-type levels. Some of the mutant RGCs even persist for some time but still have aberrant gene expression. Consistent with our findings, when apoptosis is inhibited, a much larger number of RGCs survived ^135^. These new findings suggest that additional factor(s) other than Atoh7 function in the transitional RPCs to promote them toward the RGC lineage. Whereas *Neurod1* and *Neurog*2, which are upregulated in the *Atoh7*-null retina, may be compensatory, they unlikely play major roles in the RGC lineage, as mutations of their genes lead to relatively minor RGC defects ^20,102,136^. We propose that the SoxC family of transcription factors, including Sox4 and Sox11, fulfill this role (Figure 10). This is based on previous reports that deletion of the SoxC genes leads to severely compromised RGC production ^97–99^, and our finding that they are expressed at high levels in the transitional RPCs and that their expression is not dependent on Atoh7. The SoxC factors likely also function in differentiating RGCs and other cell types, as they continue to be expressed in fate-committed retinal neurons. Thus, activation of early RGC genes such as *Pou4f2* and *Isl1*, likely requires both upstream inputs, but in the absence of Atoh7, the SoxC factors still activate some of the RGC genes (Figure 10). Ectopic expression of Pou4f2 and Isl1 together rescues RGC formation caused by deletion of *Atoh7*, and the two factors were proposed to be part of a core group of factors determining the RGC fate ^59^. In light of our current findings, the determination of the RGC fate is likely a gradual process over a time window without a clear boundary, and the function of Pou4f2 and Isl1 is to stabilize the developmental trajectory by activating genes essential for RGC differentiation and repressing genes for other fates. Some of the RGC genes are activated already by Atoh7 and/or other factors independent of Pou4f2 and Isl1, but require Pou4f2 and Isl1 to reach full amplitudes of expression, whereas many other RGC genes can only be activated by Pou4f2 and Isl1 (Figure 10). Other than activating RGC genes, Atoh7 is also involved in other aspects of RGC genesis, such as cell cycle exit, downregulation of the Notch pathway, and even generation of other cell types ^32,39,130^. Elucidating the full function of Atoh7 requires identification of it direct targets and the associated epigenetic status.

Our study also demonstrates that regular RNA-seq and scRNA-seq complement each other and can be used in combination to provide much richer information regarding transcriptomic changes due to genetic perturbations. Although regular RNA-seq lacks cellular resolution, it is more sensitive in detecting genes expressed at low levels and/or in a smaller number of cells. On the other hand, scRNA-seq enables classification of cell states/types in complex tissues and provides insights into relationships among the different cell states/types. scRNA-seq also provides precise information regarding changes in gene expression in specific cell states/types. It is worth noting that, currently, likely due to limitations of sequence depth and library construction, genes expressed at low levels and/or in small numbers of cells are not always readily detectable by scRNA-seq, but this may change as the technology further matures.

In summary, we used RNA-seq and scRNA-seq to survey gene expression in the developing retina and identify changes associated with deletion of key transcription factor genes for the RGC lineage. Our results provide a global view of the gene expression, cell states, and their relationships in early retinal development. Furthermore, our study validates and further defines a transitional state shared by all early retinal cell fates (Figure 10). Atoh7, likely in collaboration with other factors, functions within this cell state to shepherd RPCs to the RGC lineage by competing with other lineage factors and activating RGC-specific genes. Further analysis of the shifts in the epigenetic landscape along individual trajectories in both wild-type and mutant retinas will help elucidate the underlying mechanisms of RGC differentiation.

## Acknowledgments

We thank other members of the Mu laboratory and members of the Department of Ophthalmology and the Developmental Genomics Group, University of Buffalo, for helpful discussions. We also would like to thank Andrew Kelly for technical support. The knockin mouse lines were created by the Gene Targeting and Transgenic Resource Core at the Roswell Park Cancer Institute. Construction and sequencing of RNA-seq and scRNA-seq libraries were carried out at the Genomics and Bioinformatics Core of University at Buffalo. This project was supported by grants from the BrightFocus Foundation (G2016024) and the National Eye Institute of the National Institutes of Health (R01EY020545 and R01EY029705) to X.M. The content is solely the responsibility of the authors and does not necessarily represent the official views of the National Institutes of Health.

## Supplementary figure legends

**Supplementary Figure 1.**
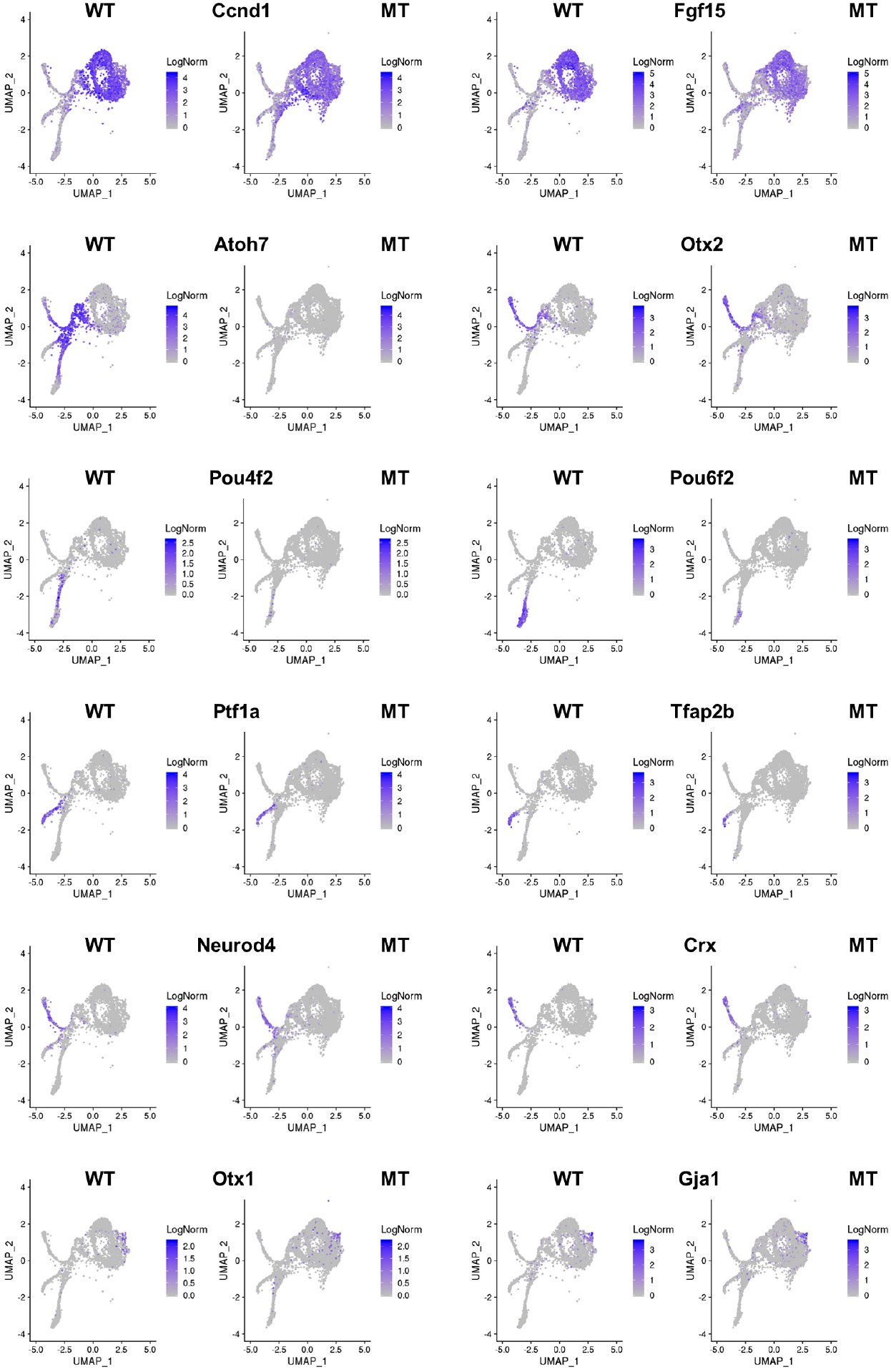
Feature plots demonstrating the cluster-specific expression of marker genes in both wild-type (WT) and *Atoh7*-null (MT) retinas at E13.5. These markers were used to assign identities to the clusters, including *Ccnd1* and *Fgf15* for naïve RPCs, *Atoh7* and *Otx2* for transitional RPCs, *Pou4f2* and *Pou6f2* for RGCs, *Ptf1a* and *Tfap2b* for amacrine and horizontal precursors, *Neurod4* and *Crx* for photoreceptors, and *Otx1* and *Gja1* for ciliary margin cells. Note that that expression of *Atoh7* and the two RGC marker genes *Pou4f2* and *Pou6f2* are diminished in the MT cells.

**Supplementary Figure 2.**
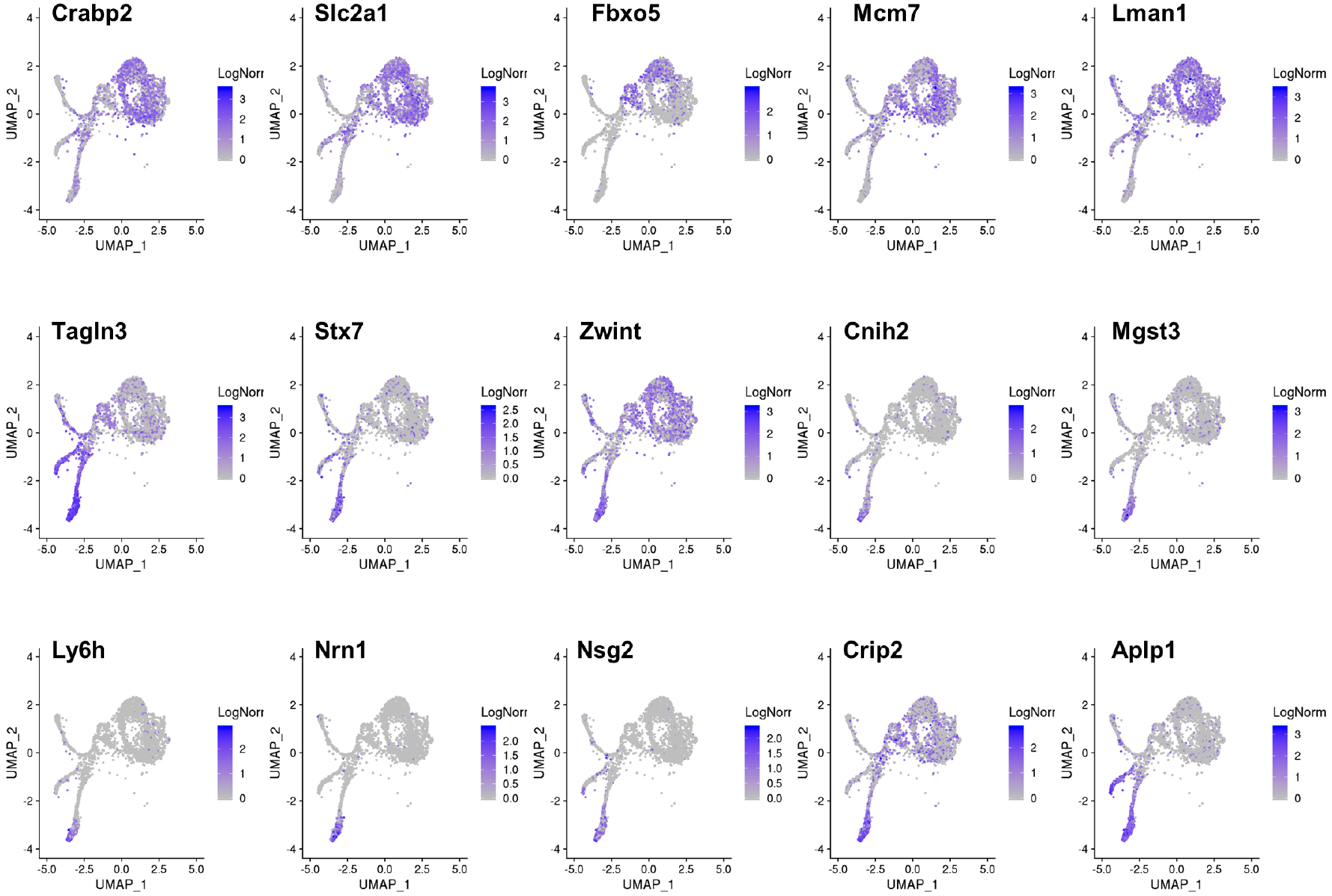
Feature plots based on scRNA-seq showing the expression patterns of five naïve RPC enriched genes and ten RGC-enriched genes at E13.5. Comparing with in situ hybridization (see Suppl. Figure 3), scRNA-seq provides more details of cell type-specific expression. For example, *Fbxo5* is expressed only in subsets of naïve RPCs and transitional RPCs, which are likely in the late S and early G2/M phases of the cell cycle (compare with Figure 3d).

**Supplementary Figure 3.**
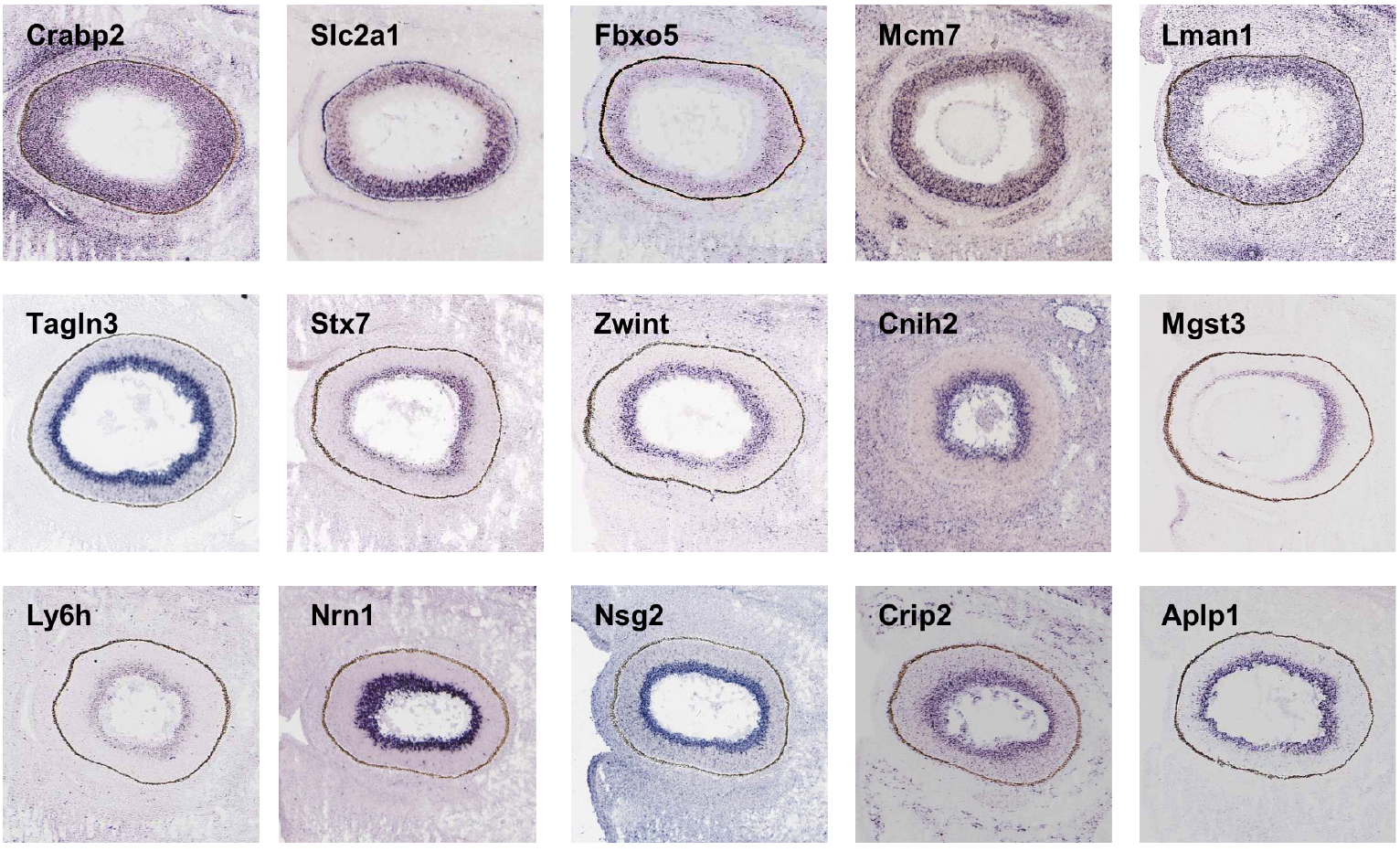
In situ hybridization confirms the spatial expression patterns of five naïve RPC enriched genes and ten RGC-enriched genes. The in situ hybridization data were obtained from the Eurexpress database and match well with the enrichment results from our scRNA-seq analysis (See Suppl. Figure 2 and Suppl. Table 9).

**Supplementary Figure 4.**
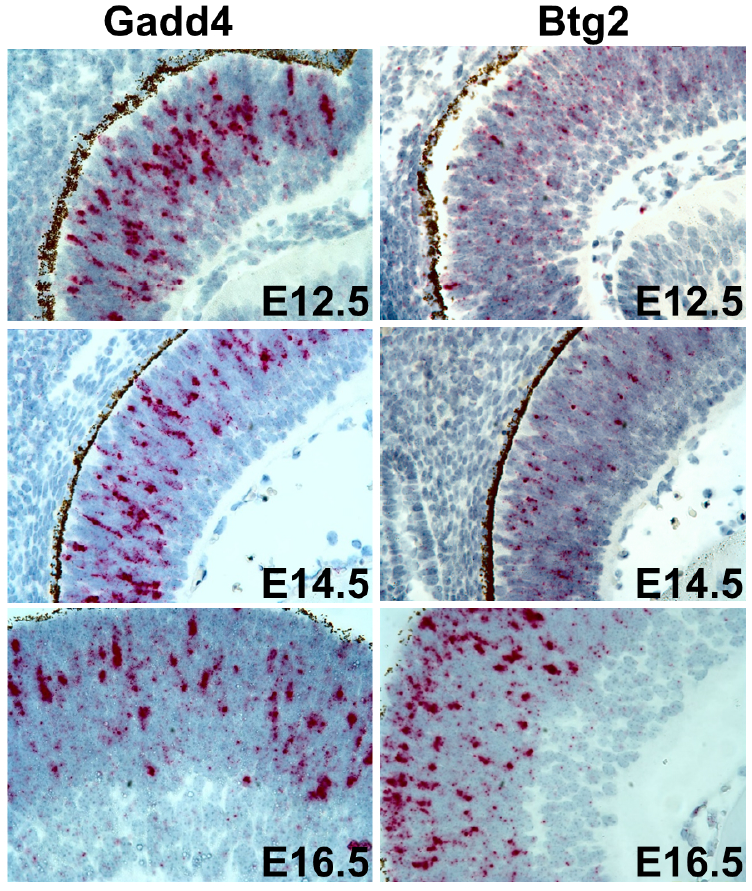
In situ hybridization with RNAscope probes confirms that *Gadd45a* and *Btg2* are expressed in subsets of RPCs with patterns similar to *Atoh7* in all three developmental stages (E12.5, E14.5, E16.5) examined.

**Supplementary Figure 5.**
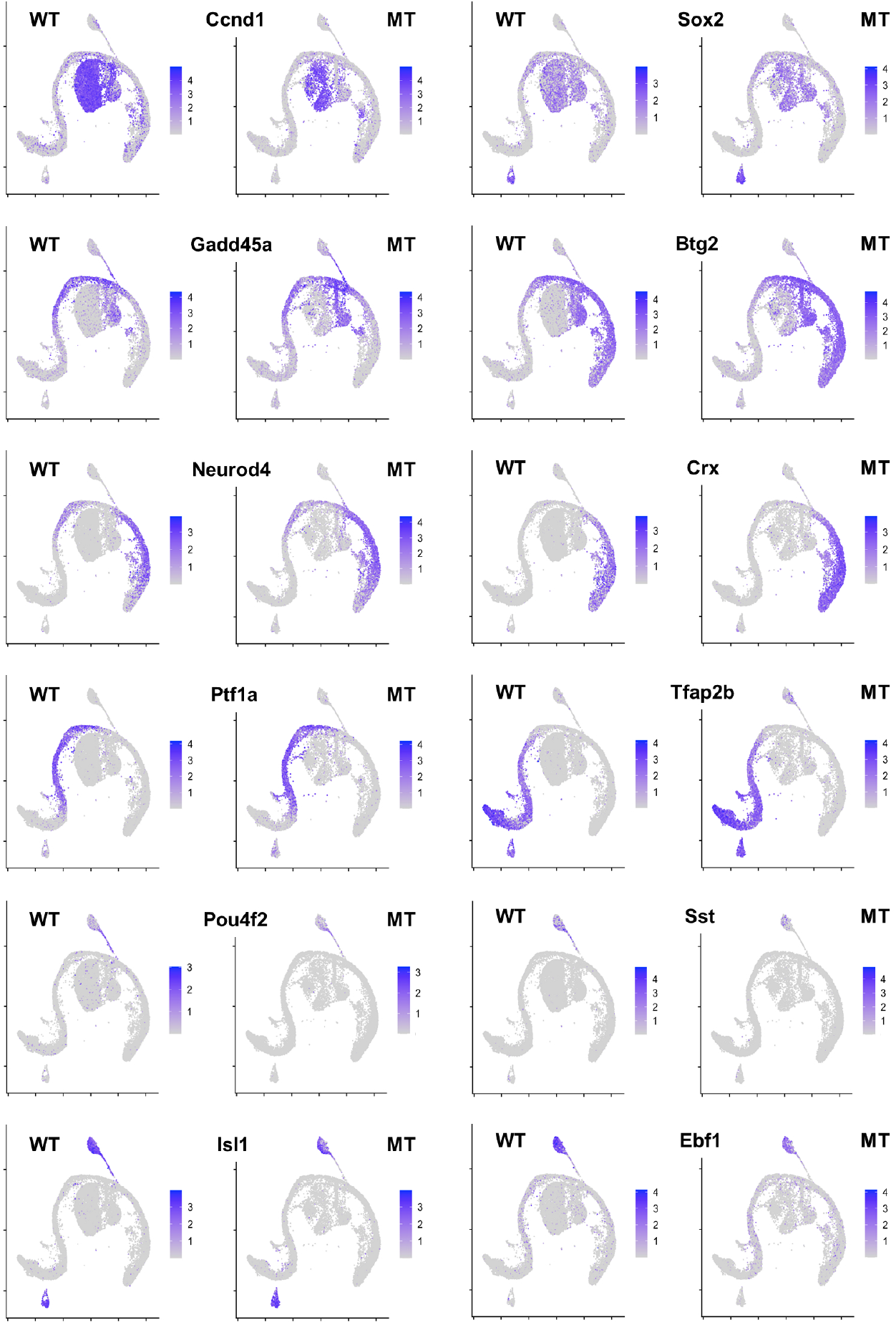
Feature plots of additional marker genes to identify clusters of the E17.5 scRNA-seq data. These markers include *Ccnd1* and *Sox2* for naïve RPCs, *Gadd45a* and *Btg2* for transitional RPCs, *Neurod4*, and *Crx* for photoreceptors, *Ptf1a* and *Tfab2b* for horizontal and amacrine cells, and *Pou4f2, Sst, Isl1*, and *Ebf1* for RGCs. WT is wild-type and MT is *Atoh7*-null.

**Supplementary Figure 6.**
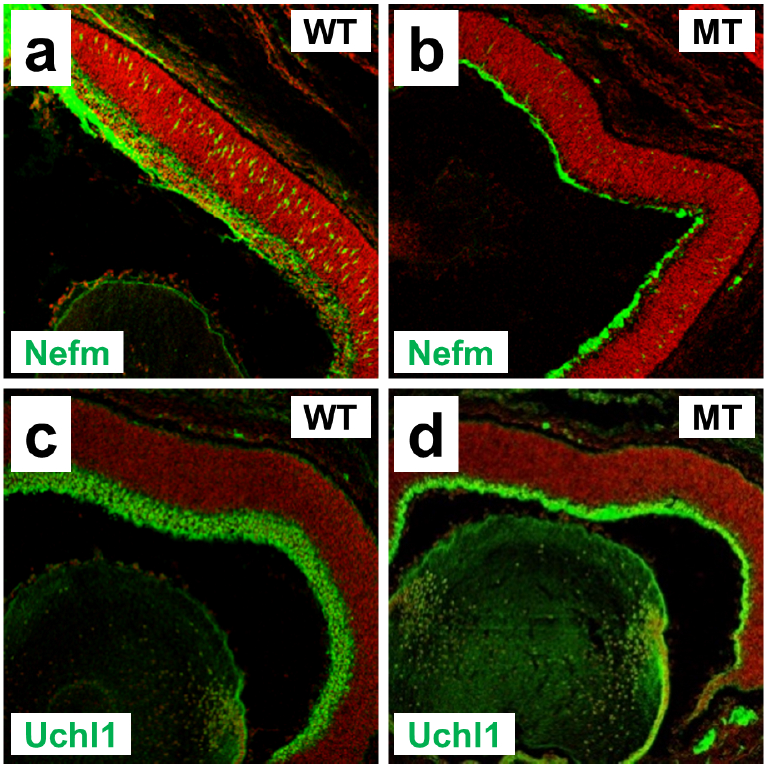
Immunofluorescence staining for two RGC markers reveals that some RGCs persist in the *Atoh7*-null retina at E17.5. a. b. Expression of neurofilament middle chain (Nefm) in wild-type (WT) and *Atoh7*-null (MT) retinas. c. d. Expression of ubiquitin carboxy-terminal hydrolase L1 (Uchl1) in WT and MT retinas. Red is counterstaining by propidium iodide.

## List of Supplementary tables

Supplementary Table 1. All genes expressed in E14.5 wild-type retina

Supplementary Table 2. Differentially expressed genes in E14.5 Atoh7-null retina

Supplementary Table 3. Differentially expressed genes in E14.5 Pou4f2-null retina

Supplementary Table 4. Differentially expressed genes in E14.5 Isl1-null retina

Supplementary Table 5. Gene ontology analysis by DAVID of DEGs in E14.5 *Atoh7*-, *Pou4f2*-, and *Isl1*-null retinas

Supplementary Table 6. Genes enriched in individual wild-type clusters

Supplementary Table 7. Stats of cluster-enriched genes

Supplementary Table 8. DEGs in C0 (270 genes)

Supplementary Table 9. DEGs in C1 (203 genes)

Supplementary Table 10. DEGs in C2 (216 genes)

Supplementary Table 11. DEGs in C3 (229 genes)

Supplementary Table 12. DEGs in C4 (128 genes)

Supplementary Table 13. DEGs in C5 (450 genes)

Supplementary Table 14. DEGs in C6 (618 genes)

Supplementary Table 15. DEGs in C7 (114 genes)

Supplementary Table 16. DEGs in C8 (58 genes)

Supplementary Table 17. DEGs in C9 (55 genes)

Supplementary Table 18. DEGs in C10 (94 genes)

Supplementary Table 19. Gene Ontology analysis of DEGs in naïve RPCs, transitional RPCs, and RGCs

Supplementary Table 20. Groups of C5/6 DEGs and enriched genes by K-means clustering

Supplementary Table 21. DEGs in E17.5 naïve Atoh7-null RPCs

Supplementary Table 22. DEGs in E17.5 transitional Atoh7-null RPCs (1)

Supplementary Table 23. DEGs in E17.5 transitional Atoh7-null RPCs (2)

Supplementary Table 24. DEGs in E17.5 Atoh7-null RGCs

## References

1. Cepko, C. Intrinsically different retinal progenitor cells produce specific types of progeny. Nat Rev Neurosci 15, 615–27 (2014).

2. Cepko, C. L., Austin, C. P., Yang, X., Alexiades, M. & Ezzeddine, D. Cell fate determination in the vertebrate retina. Proc Natl Acad Sci U A 93, 589–95 (1996).

3. Livesey, F. J. & Cepko, C. L. Vertebrate neural cell-fate determination: lessons from the retina. Nat Rev Neurosci 2, 109–18 (2001).

4. Young, R. W. Cell differentiation in the retina of the mouse. Anat Rec 212, 199–205 (1985).

5. Austin, C. P., Feldman, D. E., Ida, J. A., Jr. & Cepko, C. L. Vertebrate retinal ganglion cells are selected from competent progenitors by the action of Notch. Development 121, 3637–50 (1995).

6. Belecky-Adams, T., Cook, B. & Adler, R. Correlations between terminal mitosis and differentiated fate of retinal precursor cells in vivo and in vitro: analysis with the ‘window-labeling’ technique. Dev Biol 178, 304–15 (1996).

7. Cayouette, M., Barres, B. A. & Raff, M. Importance of intrinsic mechanisms in cell fate decisions in the developing rat retina. Neuron 40, 897–904 (2003).

8. Gomes, F. L. et al. Reconstruction of rat retinal progenitor cell lineages in vitro reveals a surprising degree of stochasticity in cell fate decisions. Development 138, 227–35 (2011).

9. La Vail, M. M., Rapaport, D. H. & Rakic, P. Cytogenesis in the monkey retina. J Comp Neurol 309, 86–114 (1991).

10. Rapaport, D. H., Wong, L. L., Wood, E. D., Yasumura, D. & LaVail, M. M. Timing and topography of cell genesis in the rat retina. J Comp Neurol 474, 304–24 (2004).

11. Reh, T. A. & Kljavin, I. J. Age of differentiation determines rat retinal germinal cell phenotype: induction of differentiation by dissociation. J Neurosci 9, 4179–89 (1989).

12. Wong, L. L. & Rapaport, D. H. Defining retinal progenitor cell competence in Xenopus laevis by clonal analysis. Development 136, 1707–15 (2009).

13. Clark, B. S. et al. Single-Cell RNA-Seq Analysis of Retinal Development Identifies NFI Factors as Regulating Mitotic Exit and Late-Born Cell Specification. Neuron 102, 1111–1126 (2019).

14. Gordon, P. J. et al. Lhx2 balances progenitor maintenance with neurogenic output and promotes competence state progression in the developing retina. J Neurosci 33, 12197–207 (2013).

15. La Torre, A., Georgi, S. & Reh, T. A. Conserved microRNA pathway regulates developmental timing of retinal neurogenesis. Proc Natl Acad Sci U A 110, E2362–70 (2013).

16. Mattar, P., Ericson, J., Blackshaw, S. & Cayouette, M. A conserved regulatory logic controls temporal identity in mouse neural progenitors. Neuron 85, 497–504 (2015).

17. Brzezinski, J. A. th, Kim, E. J., Johnson, J. E. & Reh, T. A. Ascl1 expression defines a subpopulation of lineage-restricted progenitors in the mammalian retina. Development 138, 3519–31 (2011).

18. Fu, X. et al. Epitope-tagging Math5 and Pou4f2: new tools to study retinal ganglion cell development in the mouse. Dev Dyn 238, 2309–17 (2009).

19. Hafler, B. P. et al. Transcription factor Olig2 defines subpopulations of retinal progenitor cells biased toward specific cell fates. Proc Natl Acad Sci U A 109, 7882–7 (2012).

20. Kiyama, T. et al. Overlapping spatiotemporal patterns of regulatory gene expression are required for neuronal progenitors to specify retinal ganglion cell fate. Vis. Res 51, 251–9 (2011).

21. Li, S. et al. Foxn4 controls the genesis of amacrine and horizontal cells by retinal progenitors. Neuron 43, 795–807 (2004).

22. Mu, X. et al. A gene network downstream of transcription factor Math5 regulates retinal progenitor cell competence and ganglion cell fate. Dev Biol 280, 467–81 (2005).

23. Nishida, A. et al. Otx2 homeobox gene controls retinal photoreceptor cell fate and pineal gland development. Nat Neurosci 6, 1255–63 (2003).

24. Trimarchi, J. M., Stadler, M. B. & Cepko, C. L. Individual retinal progenitor cells display extensive heterogeneity of gene expression. PLoS One 3, e1588 (2008).

25. Brown, N. L., Patel, S., Brzezinski, J. & Glaser, T. Math5 is required for retinal ganglion cell and optic nerve formation. Development 128, 2497–508 (2001).

26. Emerson, M. M., Surzenko, N., Goetz, J. J., Trimarchi, J. & Cepko, C. L. Otx2 and Onecut1 promote the fates of cone photoreceptors and horizontal cells and repress rod photoreceptors. Dev Cell 26, 59–72 (2013).

27. Fujitani, Y. et al. Ptf1a determines horizontal and amacrine cell fates during mouse retinal development. Development 133, 4439–50 (2006).

28. Nakhai, H. et al. Ptf1a is essential for the differentiation of GABAergic and glycinergic amacrine cells and horizontal cells in the mouse retina. Development 134, 1151–60 (2007).

29. Sapkota, D. et al. Onecut1 and Onecut2 redundantly regulate early retinal cell fates during development. Proc Natl Acad Sci U A 111, E4086–4095 (2014).

30. Wang, S. W. et al. Requirement for math5 in the development of retinal ganglion cells. Genes Dev 15, 24–9 (2001).

31. Brzezinski, J. A. th, Prasov, L. & Glaser, T. Math5 defines the ganglion cell competence state in a subpopulation of retinal progenitor cells exiting the cell cycle. Dev Biol 365, 395–413 (2012).

32. Feng, L. et al. MATH5 controls the acquisition of multiple retinal cell fates. Mol Brain 3, 36 (2010).

33. Bassett, E. A. & Wallace, V. A. Cell fate determination in the vertebrate retina. Trends Neurosci 35, 565–73 (2012).

34. Swaroop, A., Kim, D. & Forrest, D. Transcriptional regulation of photoreceptor development and homeostasis in the mammalian retina. Nat Rev Neurosci 11, 563–76 (2010).

35. Xiang, M. Intrinsic control of mammalian retinogenesis. Cell Mol Life Sci 70, 2519–32 (2013).

36. Macosko, E. Z. et al. Highly Parallel Genome-wide Expression Profiling of Individual Cells Using Nanoliter Droplets. Cell 161, 1202–1214 (2015).

37. Zheng, G. X. Y. et al. Massively parallel digital transcriptional profiling of single cells. Nat. Commun. 8, 1–12 (2017).

38. Giudice, Q. L., Leleu, M., Manno, G. L. & Fabre, P. J. Single-cell transcriptional logic of cell-fate specification and axon guidance in early-born retinal neurons. Development 146, dev178103 (2019).

39. Lu, Y. et al. Single-Cell Analysis of Human Retina Identifies Evolutionarily Conserved and Species-Specific Mechanisms Controlling Development. Dev. Cell (2020) doi:10.1016/j.devcel.2020.04.009.

40. Lukowski, S. W. et al. A single-cell transcriptome atlas of the adult human retina. EMBO J. e100811 (2019) doi:10.15252/embj.2018100811.

41. Menon, M. et al. Single-cell transcriptomic atlas of the human retina identifies cell types associated with age-related macular degeneration. Nat. Commun. 10, 1–9 (2019).

42. Rheaume, B. A. et al. Single cell transcriptome profiling of retinal ganglion cells identifies cellular subtypes. Nat. Commun. 9, 2759 (2018).

43. Shekhar, K. et al. Comprehensive Classification of Retinal Bipolar Neurons by Single-Cell Transcriptomics. Cell 166, 1308–1323 (2016).

44. Sridhar, A. et al. Single-Cell Transcriptomic Comparison of Human Fetal Retina, hPSC-Derived Retinal Organoids, and Long-Term Retinal Cultures. Cell Rep. 30, 1644–1659.e4 (2020).

45. Tran, N. M. et al. Single-Cell Profiles of Retinal Ganglion Cells Differing in Resilience to Injury Reveal Neuroprotective Genes. Neuron 104, 1039–1055.e12 (2019).

46. Yang, Z., Ding, K., Pan, L., Deng, M. & Gan, L. Math5 determines the competence state of retinal ganglion cell progenitors. Dev Biol 264, 240–54 (2003).

47. Erkman, L. et al. Role of transcription factors Brn-3.1 and Brn-3.2 in auditory and visual system development. Nature 381, 603–6 (1996).

48. Gan, L. et al. POU domain factor Brn-3b is required for the development of a large set of retinal ganglion cells. Proc Natl Acad Sci U A 93, 3920–5 (1996).

49. Mu, X., Fu, X., Beremand, P. D., Thomas, T. L. & Klein, W. H. Gene regulation logic in retinal ganglion cell development: Isl1 defines a critical branch distinct from but overlapping with Pou4f2. Proc Natl Acad Sci U A 105, 6942–7 (2008).

50. Pan, L., Deng, M., Xie, X. & Gan, L. ISL1 and BRN3B co-regulate the differentiation of murine retinal ganglion cells. Development 135, 1981–90 (2008).

51. Wang, S. W., Gan, L., Martin, S. E. & Klein, W. H. Abnormal polarization and axon outgrowth in retinal ganglion cells lacking the POU-domain transcription factor Brn- 3b. Mol Cell Neurosci 16, 141–56 (2000).

52. Ge, Y., Wu, F., Cheng, M., Bard, J. & Mu, X. Two new genetically modified mouse alleles labeling distinct phases of retinal ganglion cell development by fluorescent proteins. Dev. Dyn. Off. Publ. Am. Assoc. Anat. (2020) doi:10.1002/dvdy.233.

53. Gan, L., Wang, S. W., Huang, Z. & Klein, W. H. POU domain factor Brn-3b is essential for retinal ganglion cell differentiation and survival but not for initial cell fate specification. Dev Biol 210, 469–80 (1999).

54. Dobin, A. et al. STAR: ultrafast universal RNA-seq aligner. Bioinformatics 29, 15–21 (2013).

55. Robinson, M. D., McCarthy, D. J. & Smyth, G. K. edgeR: a Bioconductor package for differential expression analysis of digital gene expression data. Bioinformatics 26, 139–40 (2010).

56. Wu, F., Sapkota, D., Li, R. & Mu, X. Onecut 1 and Onecut 2 are potential regulators of mouse retinal development. J Comp Neurol 520, 952–69 (2012).

57. Tirosh, I. et al. Dissecting the multicellular ecosystem of metastatic melanoma by single-cell RNA-seq. Science 352, 189–196 (2016).

58. Haghverdi, L., Büttner, M., Wolf, F. A., Buettner, F. & Theis, F. J. Diffusion pseudotime robustly reconstructs lineage branching. Nat. Methods 13, 845–848 (2016).

59. Wu, F. et al. Two transcription factors, Pou4f2 and Isl1, are sufficient to specify the retinal ganglion cell fate. Proc Natl Acad Sci U A 112, E1559–68 (2015).

60. Mu, X. et al. Discrete gene sets depend on POU domain transcription factor Brn3b/Brn-3.2/POU4f2 for their expression in the mouse embryonic retina. Development 131, 1197–210 (2004).

61. Qiu, F., Jiang, H. & Xiang, M. A comprehensive negative regulatory program controlled by Brn3b to ensure ganglion cell specification from multipotential retinal precursors. J Neurosci 28, 3392–403 (2008).

62. Huang, D. W., Sherman, B. T. & Lempicki, R. A. Bioinformatics enrichment tools: paths toward the comprehensive functional analysis of large gene lists. Nucleic Acids Res. 37, 1–13 (2009).

63. de Melo, J. et al. Dlx1 and Dlx2 function is necessary for terminal differentiation and survival of late-born retinal ganglion cells in the developing mouse retina. Dev. Camb. Engl. 132, 311–322 (2005).

64. Zhang, Q. et al. Regulation of Brn3b by DLX1 and DLX2 is required for retinal ganglion cell differentiation in the vertebrate retina. Dev. Camb. Engl. 144, 1698–1711 (2017).

65. Mu, X. et al. Ganglion cells are required for normal progenitor- cell proliferation but not cell-fate determination or patterning in the developing mouse retina. Curr Biol 15, 525–30 (2005).

66. Sakagami, K., Gan, L. & Yang, X.-J. Distinct effects of Hedgehog signaling on neuronal fate specification and cell cycle progression in the embryonic mouse retina. J. Neurosci. Off. J. Soc. Neurosci. 29, 6932–6944 (2009).

67. Wang, Y., Dakubo, G. D., Thurig, S., Mazerolle, C. J. & Wallace, V. A. Retinal ganglion cell-derived sonic hedgehog locally controls proliferation and the timing of RGC development in the embryonic mouse retina. Development 132, 5103–13 (2005).

68. Zhang, X. M. & Yang, X. J. Regulation of retinal ganglion cell production by Sonic hedgehog. Development 128, 943–57 (2001).

69. Bosanac, I. et al. The structure of SHH in complex with HHIP reveals a recognition role for the Shh pseudo active site in signaling. Nat Struct Mol Biol 16, 691–7 (2009).

70. Lee, R. T. H., Zhao, Z. & Ingham, P. W. Hedgehog signalling. Development 143, 367–372 (2016).

71. Varjosalo, M. & Taipale, J. Hedgehog: functions and mechanisms. Genes Dev. 22, 2454–2472 (2008).

72. Kim, J. et al. GDF11 controls the timing of progenitor cell competence in developing retina. Science 308, 1927–30 (2005).

73. Hashimoto, T., Zhang, X. M., Chen, B. Y. & Yang, X. J. VEGF activates divergent intracellular signaling components to regulate retinal progenitor cell proliferation and neuronal differentiation. Development 133, 2201–10 (2006).

74. Wall, D. S. et al. Progenitor cell proliferation in the retina is dependent on Notch-independent Sonic hedgehog/Hes1 activity. J Cell Biol 184, 101–12 (2009).

75. Stuart, T. et al. Comprehensive Integration of Single-Cell Data. Cell 177, 1888–1902.e21 (2019).

76. Taranova, O. V. et al. SOX2 is a dose-dependent regulator of retinal neural progenitor competence. Genes Dev 20, 1187–202 (2006).

77. Brown, N. L. et al. Math5 encodes a murine basic helix-loop-helix transcription factor expressed during early stages of retinal neurogenesis. Development 125, 4821–33 (1998).

78. Xiang, M. et al. The Brn-3 family of POU-domain factors: primary structure, binding specificity, and expression in subsets of retinal ganglion cells and somatosensory neurons. J Neurosci 15, 4762–85 (1995).

79. Zhou, H., Yoshioka, T. & Nathans, J. Retina-derived POU-domain factor-1: a complex POU-domain gene implicated in the development of retinal ganglion and amacrine cells. J Neurosci 16, 2261–74 (1996).

80. Akagi, T. et al. Requirement of multiple basic helix-loop-helix genes for retinal neuronal subtype specification. J Biol Chem 279, 28492–8 (2004).

81. Furukawa, T., Morrow, E. M. & Cepko, C. L. Crx, a novel otx-like homeobox gene, shows photoreceptor-specific expression and regulates photoreceptor differentiation. Cell 91, 531–41 (1997).

82. Calera, M. R. et al. Connexin43 is required for production of the aqueous humor in the murine eye. J. Cell Sci. 119, 4510–4519 (2006).

83. Zhao, S., Chen, Q., Hung, F.-C. & Overbeek, P. A. BMP signaling is required for development of the ciliary body. Development 129, 4435–4442 (2002).

84. Blackshaw, S. et al. Genomic analysis of mouse retinal development. PLoS Biol 2, E247 (2004).

85. Green, E. S., Stubbs, J. L. & Levine, E. M. Genetic rescue of cell number in a mouse model of microphthalmia: interactions between Chx10 and G1-phase cell cycle regulators. Development 130, 539–52 (2003).

86. de Melo, J. et al. Lhx2 Is an Essential Factor for Retinal Gliogenesis and Notch Signaling. J Neurosci 36, 2391–405 (2016).

87. Brzezinski, J. A. th, Uoon Park, K. & Reh, T. A. Blimp1 (Prdm1) prevents re-specification of photoreceptors into retinal bipolar cells by restricting competence. Dev Biol 384, 194–204 (2013).

88. Ng, L. et al. A thyroid hormone receptor that is required for the development of green cone photoreceptors. Nat Genet 27, 94–8 (2001).

89. Rodgers, H. M., Belcastro, M., Sokolov, M. & Mathers, P. H. Embryonic markers of cone differentiation. http://www.molvis.org/molvis/v22/1455/ (2016).

90. Trimarchi, J. M. et al. Molecular heterogeneity of developing retinal ganglion and amacrine cells revealed through single cell gene expression profiling. J Comp Neurol 502, 1047–65 (2007).

91. Bassett, E. A. et al. Overlapping expression patterns and redundant roles for AP-2 transcription factors in the developing mammalian retina. Dev Dyn 241, 814–29 (2012).

92. Goodson, N. B. et al. Prdm13 is required for Ebf3+ amacrine cell formation in the retina. Dev. Biol. 434, 149–163 (2018).

93. Bélanger, M.-C., Robert, B. & Cayouette, M. Msx1-Positive Progenitors in the Retinal Ciliary Margin Give Rise to Both Neural and Non-neural Progenies in Mammals. Dev. Cell 40, 137–150 (2017).

94. Marcucci, F. et al. The Ciliary Margin Zone of the Mammalian Retina Generates Retinal Ganglion Cells. Cell Rep. 17, 3153–3164 (2016).

95. Pennesi, M. E. et al. BETA2/NeuroD1 null mice: a new model for transcription factor-dependent photoreceptor degeneration. J Neurosci 23, 453–61 (2003).

96. Diez-Roux, G. et al. A High-Resolution Anatomical Atlas of the Transcriptome in the Mouse Embryo. PLoS Biol. 9, (2011).

97. Chang, K.-C. et al. Novel Regulatory Mechanisms for the SoxC Transcriptional Network Required for Visual Pathway Development. J. Neurosci. Off. J. Soc. Neurosci. 37, 4967–4981 (2017).

98. Jiang, Y. et al. Transcription factors SOX4 and SOX11 function redundantly to regulate the development of mouse retinal ganglion cells. J Biol Chem 288, 18429–38 (2013).

99. Kuwajima, T., Soares, C. A., Sitko, A. A., Lefebvre, V. & Mason, C. SoxC Transcription Factors Promote Contralateral Retinal Ganglion Cell Differentiation and Axon Guidance in the Mouse Visual System. Neuron 93, 1110–1125.e5 (2017).

100. Usui, A. et al. The early retinal progenitor-expressed gene Sox11 regulates the timing of the differentiation of retinal cells. Development 140, 740–750 (2013).

101. Wang, F. et al. RNAscope: a novel in situ RNA analysis platform for formalin-fixed, paraffin-embedded tissues. J Mol Diagn 14, 22–9 (2012).

102. Hufnagel, R. B., Le, T. T., Riesenberg, A. L. & Brown, N. L. Neurog2 controls the leading edge of neurogenesis in the mammalian retina. Dev Biol 340, 490–503 (2010).

103. Feng, L. et al. Requirement for Bhlhb5 in the specification of amacrine and cone bipolar subtypes in mouse retina. Development 133, 4815–25 (2006).

104. Feng, L. et al. Brn-3b inhibits generation of amacrine cells by binding to and negatively regulating DLX1/2 in developing retina. Neuroscience 195, 9–20 (2011).

105. Huang, L. et al. Bhlhb5 is required for the subtype development of retinal amacrine and bipolar cells in mice. Dev. Dyn. Off. Publ. Am. Assoc. Anat. 243, 279–289 (2014).

106. Riesenberg, A. N., Conley, K. W., Le, T. T. & Brown, N. L. Separate and coincident expression of Hes1 and Hes5 in the developing mouse eye. Dev. Dyn. Off. Publ. Am. Assoc. Anat. 247, 212–221 (2018).

107. Mao, B., Zhang, Z. & Wang, G. BTG2: A rising star of tumor suppressors (Review). Int. J. Oncol. 46, 459–464 (2015).

108. Salvador, J. M., Brown-Clay, J. D. & Fornace, A. J. Gadd45 in stress signaling, cell cycle control, and apoptosis. Adv. Exp. Med. Biol. 793, 1–19 (2013).

109. Tamura, R. E. et al. GADD45 proteins: central players in tumorigenesis. Curr. Mol. Med. 12, 634–651 (2012).

110. Yuniati, L., Scheijen, B., van der Meer, L. T. & van Leeuwen, F. N. Tumor suppressors BTG1 and BTG2: Beyond growth control. J. Cell. Physiol. 234, 5379–5389 (2019).

111. Miesfeld, J. B., Glaser, T. & Brown, N. L. The dynamics of native Atoh7 protein expression during mouse retinal histogenesis, revealed with a new antibody. Gene Expr. Patterns GEP 27, 114–121 (2018).

112. Le, T. T., Wroblewski, E., Patel, S., Riesenberg, A. N. & Brown, N. L. Math5 is required for both early retinal neuron differentiation and cell cycle progression. Dev Biol 295, 764–78 (2006).

113. Jadhav, A. P., Cho, S. H. & Cepko, C. L. Notch activity permits retinal cells to progress through multiple progenitor states and acquire a stem cell property. Proc Natl Acad Sci U A 103, 18998–9003 (2006).

114. Maurer, K. A., Riesenberg, A. N. & Brown, N. L. Notch signaling differentially regulates Atoh7 and Neurog2 in the distal mouse retina. Development 141, 3243–54 (2014).

115. Perron, M. & Harris, W. A. Determination of vertebrate retinal progenitor cell fate by the Notch pathway and basic helix-loop-helix transcription factors. Cell Mol Life Sci 57, 215–23 (2000).

116. Riesenberg, A. N. & Brown, N. L. Cell autonomous and nonautonomous requirements for Delltalike1 during early mouse retinal neurogenesis. Dev. Dyn. Off. Publ. Am. Assoc. Anat. 245, 631–640 (2016).

117. Riesenberg, A. N., Liu, Z., Kopan, R. & Brown, N. L. Rbpj cell autonomous regulation of retinal ganglion cell and cone photoreceptor fates in the mouse retina. J Neurosci 29, 12865–77 (2009).

118. Yaron, O., Farhy, C., Marquardt, T., Applebury, M. & Ashery-Padan, R. Notch1 functions to suppress cone-photoreceptor fate specification in the developing mouse retina. Development 133, 1367–78 (2006).

119. Furukawa, T., Mukherjee, S., Bao, Z. Z., Morrow, E. M. & Cepko, C. L. rax, Hes1, and notch1 promote the formation of Muller glia by postnatal retinal progenitor cells. Neuron 26, 383–94 (2000).

120. Lee, H. Y. et al. Multiple requirements for Hes 1 during early eye formation. Dev Biol 284, 464–78 (2005).

121. Nelson, B. R., Gumuscu, B., Hartman, B. H. & Reh, T. A. Notch activity is downregulated just prior to retinal ganglion cell differentiation. Dev Neurosci 28, 128–41 (2006).

122. Nelson, B. R., Hartman, B. H., Georgi, S. A., Lan, M. S. & Reh, T. A. Transient inactivation of Notch signaling synchronizes differentiation of neural progenitor cells. Dev. Biol. 304, 479–498 (2007).

123. Cau, E. & Blader, P. Notch activity in the nervous system: to switch or not switch? Neural Develop. 4, 36 (2009).

124. Kageyama, R., Ohtsuka, T., Shimojo, H. & Imayoshi, I. Dynamic Notch signaling in neural progenitor cells and a revised view of lateral inhibition. Nat. Neurosci. 11, 1247–1251 (2008).

125. Lathia, J. D., Mattson, M. P. & Cheng, A. Notch: From Neural Development to Neurological Disorders. J. Neurochem. 107, 1471–1481 (2008).

126. Luo, H. et al. Forkhead box N4 (Foxn4) activates Dll4-Notch signaling to suppress photoreceptor cell fates of early retinal progenitors. Proc. Natl. Acad. Sci. U. S. A. 109, E553–562 (2012).

127. Nelson, B. R. et al. Acheate-scute like 1 (Ascl1) is required for normal delta-like (Dll) gene expression and notch signaling during retinal development. Dev. Dyn. Off. Publ. Am. Assoc. Anat. 238, 2163–2178 (2009).

128. Liu, W., Mo, Z. & Xiang, M. The Ath5 proneural genes function upstream of Brn3 POU domain transcription factor genes to promote retinal ganglion cell development. Proc Natl Acad Sci U A 98, 1649–54 (2001).

129. Prasov, L., Nagy, M., Rudolph, D. D. & Glaser, T. Math5 (Atoh7) gene dosage limits retinal ganglion cell genesis. Neuroreport 23, 631–4 (2012).

130. Zhang, X.-M., Hashimoto, T., Tang, R. & Yang, X.-J. Elevated expression of human bHLH factor ATOH7 accelerates cell cycle progression of progenitors and enhances production of avian retinal ganglion cells. Sci. Rep. 8, 6823 (2018).

131. Ali, F. et al. Cell cycle-regulated multi-site phosphorylation of Neurogenin 2 coordinates cell cycling with differentiation during neurogenesis. Dev. Camb. Engl. 138, 4267–4277 (2011).

132. Moore, K. B., Schneider, M. L. & Vetter, M. L. Posttranslational mechanisms control the timing of bHLH function and regulate retinal cell fate. Neuron 34, 183–195 (2002).

133. Satou, Y. et al. Phosphorylation states change Otx2 activity for cell proliferation and patterning in the Xenopus embryo. Dev. Camb. Engl. 145, (2018).

134. Tomic, G. et al. Phospho-regulation of ATOH1 Is Required for Plasticity of Secretory Progenitors and Tissue Regeneration. Cell Stem Cell 23, 436–443.e7 (2018).

135. Brodie-Kommit, J. et al. Atoh7-independent specification of retinal ganglion cell identity. bioRxiv 2020.05.27.116954 (2020) doi:10.1101/2020.05.27.116954.

136. Mao, C. A., Wang, S. W., Pan, P. & Klein, W. H. Rewiring the retinal ganglion cell gene regulatory network: Neurod1 promotes retinal ganglion cell fate in the absence of Math5. Development 135, 3379–88 (2008).

